# PHACTn enables training-free, context-independent inference of nucleotide variant tolerance across the genome

**DOI:** 10.64898/2026.09.08.750126

**Authors:** Ceren Yildirim, Nurdan Kuru, Ogün Adebali

## Abstract

Accurate prediction of single-nucleotide variant (SNV) tolerability across the entire human genome remains a fundamental challenge in computational genomics, particularly for non-coding regions where the regulatory landscape is vast and poorly understood. Machine learning classifiers suffer from data circularity and demographic bias, while genomic language models demand massive computational resources and offer little biological interpretability. Here, we present PHACTn (Phylogeny-Aware Computing of Tolerance for nucleotide variants), a training-free, parameter-minimal method that infers nucleotide variant tolerability by traversing the mammalian phylogenetic tree and explicitly modeling the evolutionary independence of observed substitutions and their distance from the query species. With only 4 interpretable parameters, no training and no GPU requirement, PHACTn outperforms all evaluated tools on non-coding variants curated from both the ClinVar, and on non-coding variants potentially responsible for selected Mendelian diseases curated from OMIM. Additionally, it achieves state-of-the-art performance on variants within the informative range of alignment-based inference. These results establish that principled probabilistic phylogenetic modeling captures evolutionary constraint signals that large-scale sequence models fail to recover, offering a powerful, accessible, and mechanistically transparent alternative for genome-wide variant effect prediction.

## 1 Introduction

The human genome spans approximately 3.1 billion base pairs, yet only around 2% of these encode proteins [1, 2]. Large-scale functional genomics and genome-wide association studies (GWAS) have revealed that the majority of single-nucleotide polymorphisms (SNPs) associated with diseases are clustered in remaining non-coding regulatory landscape [3, 4]. Consequently, single-nucleotide variants (SNVs) within these regions have an essential role in shaping genetic diversity, influencing both phenotypic traits and the carrier’s susceptibility to certain diseases [4, 5].

Currently, the ability to detect new SNVs has significantly improved with the affordability and advancement of high-throughput sequencing techniques. Even though sequencing identifies these variants, it is still necessary to effectively reveal their biological complexity and consequences, thereby contributing to clinical insights into genetic diseases. In particular, if the consequences of SNVs are clearly defined, the diagnosis of Mendelian diseases and the identification of the underlying causes of complex diseases can be facilitated [6]. Analysis of SNVs also serves as a foundation for precision medicine [7], evolutionary biology, and functional genomics. Since validating millions of identified variants per genome is impractical and burdensome, the field has progressively shifted towards computational methodologies [8, 9].

Machine learning (ML) classifiers have become a standard approach for predicting the effects of SNVs. These classifiers are trained on known pathogenic and benign variants combined with biological features such as conservation, amino-acid properties, structural annotations, and regulatory patterns [9–17]. However, they face significant challenges and limitations. While they benefit from labeled variants from ClinVar, HGMD, and other curated databases during training, observed variant availability is not uniform; they are biased towards particular genes, variant types, and mechanisms, and are prone to demographic imbalances due to sampling bias [18–21]. These methods are also often affected by data circularity, in which the same variants appear in both training and testing sets due to shared data sources, a problem that is particularly pronounced in ensemble meta-predictors, because they aggregate outputs from multiple individual tools [22]. As a result, ML classifiers lead to overly optimistic performance estimates that may not reflect true utility in real clinical environments [23, 24].

Genomic language models (gLMs) offer a more flexible alternative, thus having significant potential for predicting how genetic variants affect biological function without the need for manually prepared features. Unlike ML classifiers, they take raw input and learn the context of the genome, such as repetitive patterns, splice signals, regulatory grammar, and long-range dependencies directly from data. [25–27]. Mostly, the genomic language is learned through Masked Language Modeling (MLM) by hiding the nucleotide from the model during training [28]. Once the model has captured a general representation of the genome and assigned a probability to every possible nucleotide at each position, reflecting how well each base fits its surrounding sequence context, these probabilities can be directly leveraged for variant effect prediction in a zero-shot manner, without any further training, by comparing the likelihood of the reference and alternative alleles at a given position [28]. However, the application of genomic language models in clinical settings remains limited by their black-box nature, which hinders biological interpretability, and by the substantial computational resources they require.

Moreover, conventional sequence-based gLMs have demonstrated lower performance compared to alignment-based techniques in predicting causal non-coding variants, underscoring the importance of explicitly integrating evolutionary information from multispecies alignments [29]. Indeed, the history of evolution serves as a natural experiment, illustrating which genetic variations are selected for and which are eliminated. If a genomic position is naturally conserved for millions of years due to its critical function, that position is unlikely to tolerate mutations, thereby increasing the likelihood that any mutation at this site will be pathogenic. On the other hand, variable sites are less likely to harbor deleterious mutations. Thus, by understanding the conservation level of a given position, we can infer its mutation risk. PhyloP [30], PhastCons [31], and GERP [32] are statistical frameworks designed to understand evolutionary pressures on the human genome derived from multispecies whole-genome alignments; for this reason, they are still widely used as standalone metrics or features for other prediction models.

GPN-MSA [33] was recently developed as a whole-genome alignment-based genomic language model, specifically trained for variant effect prediction. Even though the model learns from multi-species alignments, relying only on that biological information presents a considerable challenge: the model cannot distinguish between nucleotide changes at the same position that arose independently versus those inherited from a common ancestor. Without considering species’ evolutionary history, treating all sequences as equally distant would lead to a false interpretation of the tolerability of a given position [34]. Therefore, integrating whole-genome alignments with ancestral trees would help to better estimate divergence times and substitution patterns.

In order to overcome these limitations, here we present a novel, training-free, phylogeny-aware probabilistic method, PHACTn, for predicting the tolerability of single-nucleotide variants.

It employs reconstructed nucleotide probabilities at each tree node to infer substitution tolerance, explicitly accounting for the evolutionary independence of observed substitutions and their phylogenetic distance from the query species.

The underlying principle of this approach, PHACT (Phylogeny-Aware Computing of Tolerance) [35], originally designed for missense mutations, has demonstrated superior predictive performance compared to widely used tools in its field. Moreover, PHACTboost (Phylogeny-Aware Pathogenicity Predictor for Missense Mutations via Boosting) [36] builds upon PHACT’s foundation by integrating gradient boosting decision trees along with tree- and alignment-related features. By outperforming state-of-the-art predictors benchmarked in dbNSFP [36], PHACTboost further validates the effectiveness of phylogeny-aware approaches for missense variant pathogenicity prediction.

Because of its success, we adapted the foundational logic of PHACT to single-nucleotide variants, while enhancing it by explicitly handling alignment gaps with gap-aware ancestral reconstruction and a weighted gap-frequency score that models indels across the species. Operating on a single position at a time without any sequence context, PHACTn achieves performance superior to conservation-based methods and competitive with alignment-based genomic language models and large-scale sequence model, Evo2, without any learned parameters or training data. PHACTn provides a more accurate and biologically meaningful interpretation, and unlike black-box models, its outputs are traceable and benefit from the complete evolutionary history of the sequences.

## 2 Results

### 2.1 Algorithm of PHACTn: Phylogeny-Aware Computing of Tolerance for Single Nucleotide Variants

Conservation scores derived only from multiple sequence alignments neglect the evolutionary relationships among substitution events, treating co-inherited changes as independent occurrences. To illustrate, two alignments in Figure 1a are identical when taking into account the observed frequencies of Thymine (T) and Cytosine (C) alone, without considering the context of the phylogenetic tree. However, given the evolutionary relationships, the recurring, independent characteristics of the first scenario provide stronger evidence for the neutrality of the C-to-T substitution. On the other hand, the second scenario demonstrates that dependent events arise from a single mutation in a common ancestor, accounting for all instances of T. This distinction has direct consequences for tolerability assessment: a single substitution event should not be overcounted by the number of its observed descendants. As a single event might have been compensated for by another change in the species where that substitution occurs, it is plausible that this change is not tolerated in the other clades of the tree. On the other hand, independent substitutions at the same position strongly suggest neutrality. Nevertheless, when evaluated solely using MSA, these significant independent alterations may not be well differentiated from dependent ones, hence underestimating their contribution to tolerability assessment. PHACTn resolves this issue by traversing the tree and calculating the probability difference across all interconnected nodes. This approach was adapted from the original PHACT’s [35] logic.

**Fig. 1:**
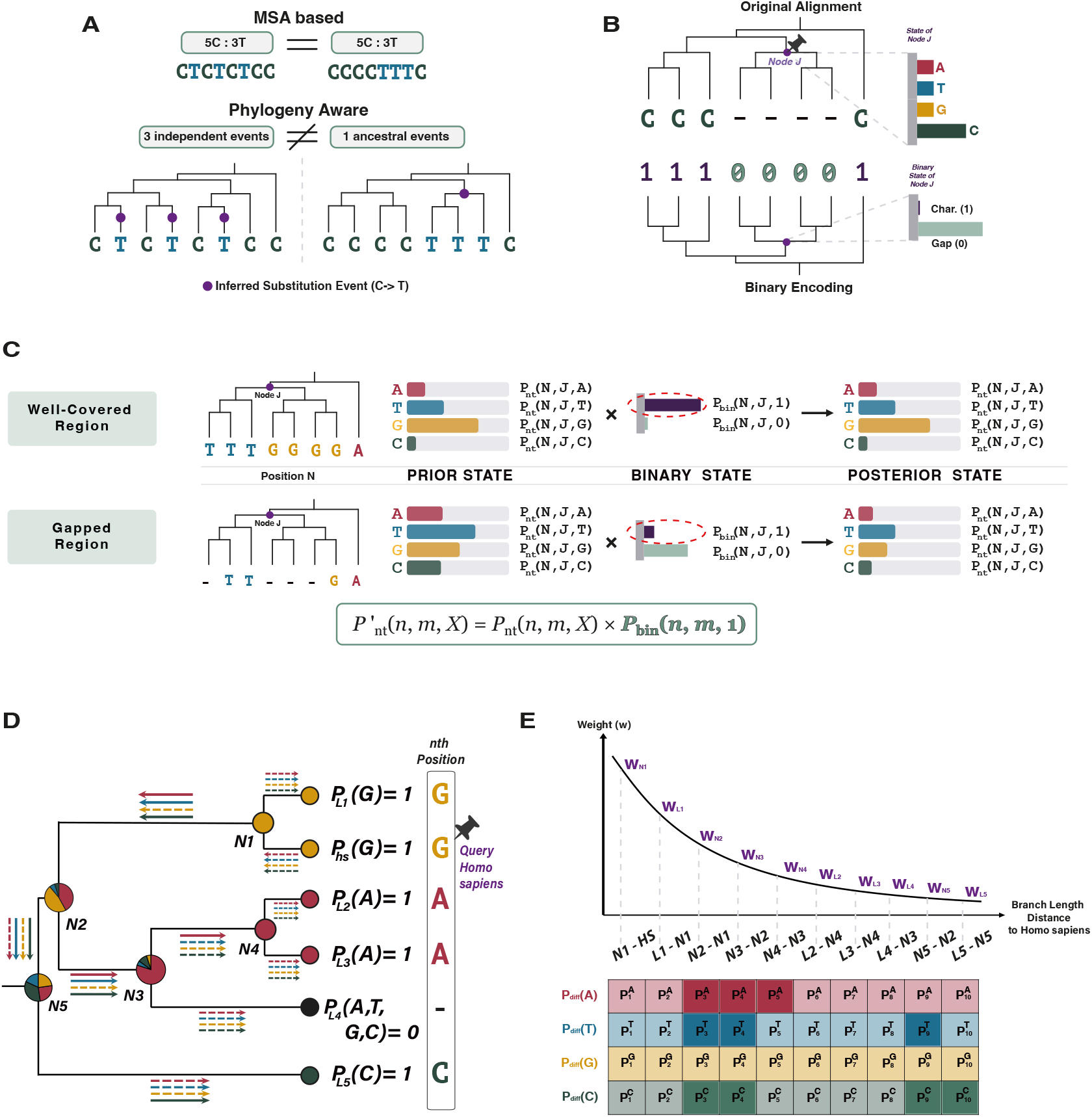
Overview of the PHACTn framework and its phylogeny-aware scoring logic. Panel descriptions are provided on the following page. **(a)** The limitation of MSA-based approaches. While they treat two alignments as equivalent because the nucleotide frequencies are identical, a phylogeny-aware point of view distinguishes independent substitution events from ancestral events. **(b)** The ancestral reconstruction tends to assign probabilities to unseen nucleotides, as in probabilities of the Node J in the original alignment. To overcome this limitation, the alignment is encoded as two states: 1 for a character (A, T, G, C) and 0 for gaps. Now, since the gap state is modeled separately, the probabilities are more accurate in the gap regions. **(c)** A method for calculating posterior probabilities. Since the probability of a gap state remains low in well-covered regions, the weighted probabilities remain essentially unchanged when multiplied by 1 gap probability, which is the character probability. However, if a region provides abundant gap information, the weighted probabilities remain low even when the assigned nucleotide probabilities are high because 1 gap probability is small. This effectively filters out noise. **(d)** Illustration of the PHACTn tree traversal logic. Starting from the query species, probability differences are computed between each pair of consecutive nodes and leaves across the entire phylogeny. Arrow colors denote the four nucleotides: red for A, yellow for G, blue for T, and green for C. Solid arrows indicate positive probability differences, corresponding to independent substitution events that contribute to the tolerance score, while dashed arrows indicate negative or zero differences, representing the propagation of previously counted events and are excluded from the summation. **(e)** Distance-dependent weighting and probability difference matrix in PHACTn. (Top) Each node and leaf is assigned a weight inversely proportional to its cumulative branch length distance to Homo sapiens, so that evolutionary events in closely related lineages contribute more to the final tolerance score. (Bottom) The matrix of positive probability differences for each nucleotide (A, T, G, C) across all nodes and leaves, where darker shading indicates larger contributions to the raw tolerance score after multiplication by the corresponding weight. For example, positive probability differences for adenine at nodes N2-N1, N3-N2, and N4-N3 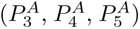 are multiplied by *W_N_*_2_, *W_N_*_3_, and *W_N_*_4_, whereas guanine yields no positive differences and contributes a score of zero. The final score is obtained by summing all weighted differences and normalizing.

For a thorough evaluation of evolutionary evidence, exploiting phylogenetic trees is known to be crucial. It has been proposed that a mutation is more likely to be benign if it is observed in species closely related to the query organism, and less likely if it occurs only in distant lineages [35, 37, 38]. Thus, PHACTn allocates a weight to each node that is inversely proportional to its phylogenetic distance from the query sequence, thereby ensuring that closely related species exert a greater influence on the final tolerance score. Despite human variants being used in this study, the algorithm places no inherent restriction on the choice of query species.

PHACTn works on nucleotide probability distributions, allowing a more continuous and statistically precise interpretation of evolutionary signals. The probabilities at terminal nodes are determined based on the nucleotide observed directly in the MSA, while ancestral reconstruction (ASR) is performed to obtain the probabilities at internal nodes. This step serves as a crucial source of information for the PHACT-like algorithms [35, 36, 39].

However, there is a fundamental limitation in the nucleotide ASR; it is blind to gaps. Standard substitution models are built for only four states: A, C, T, and G. To determine the probability of change among those states, widely used tools such as IQ-TREE [40] and RAxML-NG [41] treat gaps (-) as missing data. This eliminates the possibility that the ancestral state was a deletion, since any site must sum to 1.0 across the four nucleotides (Figure 1b, top). This limitation creates a significant problem: if a deletion is a predominant event at a specific site, nucleotide ASR may be forced to assign ghost probabilities to unseen nucleotides due to the evolutionary model’s nature. When ghost probabilities are included in PHACTn, they can accumulate, leading to misleading tolerability signals. Consequently, the algorithm may incorrectly infer an increased probability of a specific character on a branch when the sequence is actually undergoing a substitution to a gap [39]. To overcome this limitation, the binary states were also calculated (Figure 1b, bottom). Then, to obtain more accurate nucleotide states distributions, ancestral probabilities were weighted in a gap-aware manner. Thus, in cases where a position is likely to be a gap in the ancestor, when character probability is low, all nucleotide probabilities decrease proportionally, whereas in well-covered positions, the prior probabilities are largely preserved (Figure 1c). This approach has been tested in the Phylogeny-Aware Detection of Molecular Coevolution (PHACE) [39] and was shown to improve overall performance.

With gap-corrected probabilities established at each node, the next step is to traverse the tree and quantify the evolutionary change of each nucleotide relative to the query species by taking the difference in its probability between adjacent nodes to derive a tolerability score. As shown in Figure 1d, the probabilities at the nodes are represented as pie charts. The paths from *Homo sapiens* to all other nodes are indicated by four arrows corresponding to the four nucleotides; their direction is critical because it represents both the tree traversal and the probability calculation direction. For example, when moving from N1 to N2, the probability of G (represented by yellow) decreases, shown as a dashed arrow, while the remaining three nucleotides increase, shown as solid arrows; positive differences indicate a substitution event on that branch, while negative ones represent the propagation of a previously counted event through the rest of the tree. Including negative differences would lead to double-counting of dependent substitution events, so only positive ones are carried forward, weighted inversely by distance from the query and summed (Figure 1e) to generate the raw tolerability score for each nucleotide. Multiple weighting functions were tested; details will be discussed in the next section.

Although gap-aware weighted probabilities correct nucleotide distributions at the node level, this process does not reflect the evolutionary dispensability of a position in the final score. Therefore, assuming that positions where gaps have accumulated predominantly in lineages close to the query are not under strong purifying selection, an additional gap correction term has been included in the PHACTn score. The comparative performance of the baseline PHACTn score and the gap-corrected PHACTn score is discussed in the next section; however, in general, the gap-corrected formulation demonstrated better prediction performance and revealed that position-level gap information provides a meaningful additional signal beyond node-level correction.

### 2.2 Performance Evaluation

To compute genome-wide single-nucleotide variant tolerability scores, we used the UCSC 470-way mammalian whole-genome alignment [42] and its pre-built phylogenetic tree, as described in the Methods section.

Instead of using the 100-species deep-vertebrate alignment, we proceeded with the mammalian version because the greater number of species is more likely to provide better information for identifying independent evolutionary events. Also, mammalian-only comparisons provide the specificity needed to resolve functional elements relevant to human biology [43], whereas the inclusion of more distantly related vertebrates leads to a loss of specificity that can obscure elements underlying specialized mammalian-specific functions and miss critical lineage-specific signals [43, 44]. This choice is further supported by the original PHACT study, which found that tolerability scores are predominantly driven by the closest species [35].

For the primary benchmark, we constructed a variant set (details in the Methods section) using ClinVar [45] pathogenic and gnomAD [46] common variants (AF *>* 0.01).

#### 2.2.1 Gap Correction and Phylogenetic Weighting Substantially Improve PHACTn Performance

During the PHACTn score accumulation step, several approaches were tested to evaluate how different weighting schemes affect overall performance. These weight functions were built to give higher weights to species that are more closely related to the query. They were taken directly from the original PHACT [35, 36] with no alterations because they are compatible across varying topologies.

The mathematical functions and their plots of eight weighting schemes can be seen in Figure S1. They fall into two broad categories depending on the information they use: *distance-only* weights, which rely solely on branch lengths, and *dual-metric* weights, which take both branch length and the number of ancestral nodes between a given node and the query leaf into account. All possible combinations of weighting function and normalisation method were evaluated, and the resulting AUROC and AUPR values are summarised in Figure S2. The variation in performance across different weights confirms that the choice of weighting scheme plays a significant role. Dual-metric_4_ and Gaussian*_σ_*_=0.1_ achieved the highest performance among all weighting functions, reflecting that closely related species are more informative, as its weights decay steeply with both branch length and topological distance.

When comparing normalization strategies, scores normalized by the observed frequency of the query nucleotide (PHACTn-FN) outperformed those normalized by the total number of nodes (PHACTn-NN) (Figure 2a, Supplementary Figure S2). Since the observed nucleotide frequency captures whether a position is variable, this difference indicates that the diversity signal at the position level carries meaningful predictive information.

**Fig. 2:**
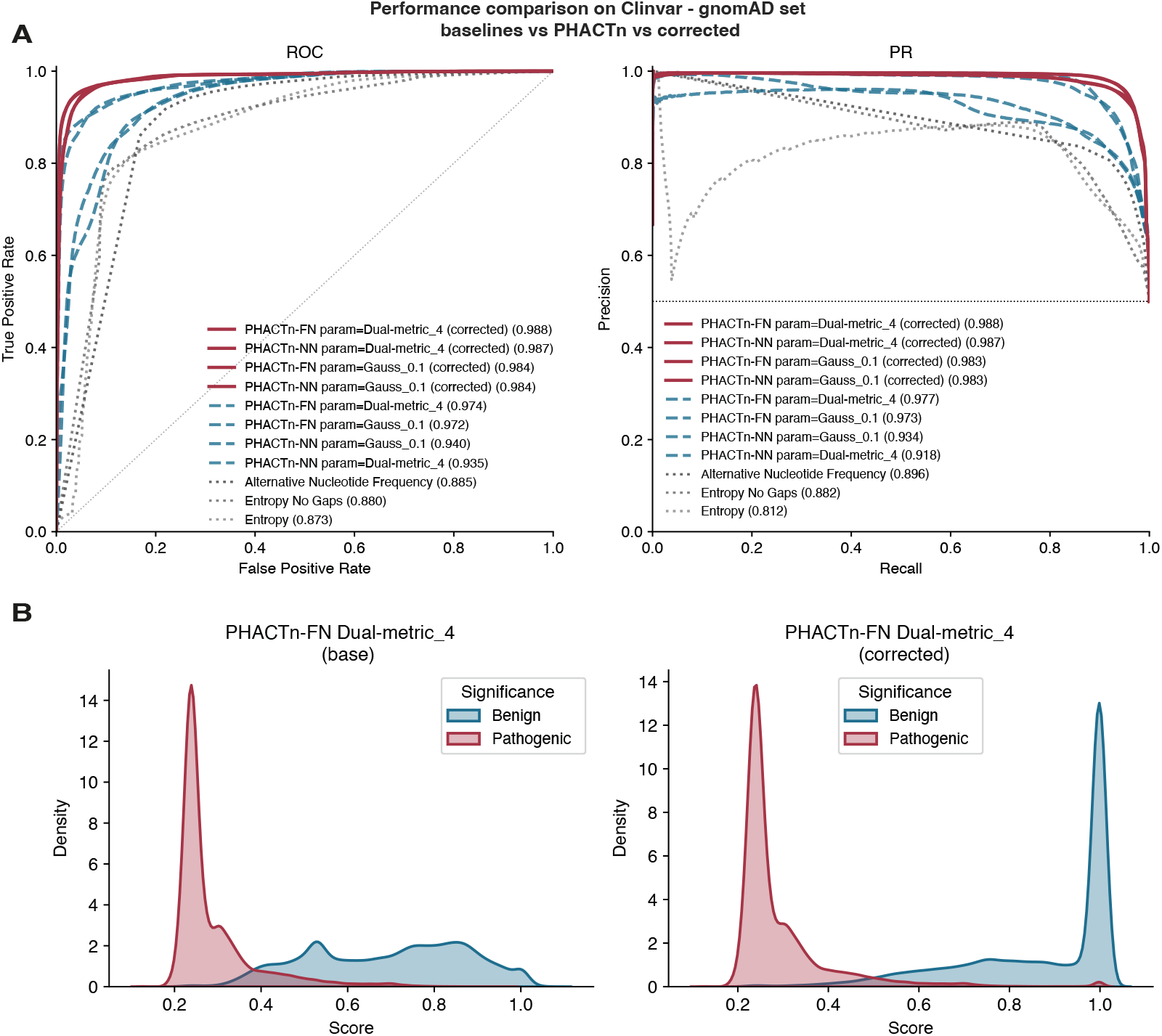
**(a)** ROC (left) and PR (right) curves of base PHACTn (blue), gap-corrected PHACTn (red), and MSA-based metrics (gray) on Clinvar-gnomAD variant set (*∼*154k benign and *∼*154k pathogenic). AUROC and AUPRC values are shown in order. **(b)** PHACTn-FN Dual-metric_4_ base and gap corrected score distribution on the ground truth labels. Red indicates pathogenic, blue indicates benign.

Gap correction further improved performance across all scores. The distribution of base and gap-corrected scores (Dual-metric_4_, Gaussian*_σ_*_=0.1_, and Gaussian*_σ_*_=0.5_) across pathogenic and benign variants is presented in Figure 2b and Supplementary Figure S3. It is observed that scores with gap correction applied to all weight functions showed more distinct separation between the two classes.

Among MSA-based baseline metrics, alternative nucleotide frequency (AUROC = 0.885, AUPR = 0.896), entropy (AUROC = 0.873, AUPR = 0.812), and gap-free entropy (AUROC = 0.880, AUPR = 0.882) all yielded substantially lower values (Figure 2a). The clear superiority of the phylogeny-aware approach, PHACTn (AUROC = 0.988, AUPR = 0.988), over all alignment-based metrics demonstrates that phylogenetic tree traversal and the evolutionary independence of gaps provide a meaningful signal beyond what alignment alone can capture.

Because of its highest performance, especially with the gap correction, we picked Dual-metric_4_ which is normalized by the observed frequency of the query nucleotide as primary PHACTn score.

#### 2.2.2 PHACTn Achieves Competitive Performance Overall and Outperforms Benchmark Tools in Non-Coding Variant Prediction

We present a performance comparison of PHACTn evaluated on the CG dataset against benchmark tools across five metrics: AUROC, AUPR (Figure 3a), F1 score, balanced accuracy, and MCC (Table 1).

**Fig. 3:**
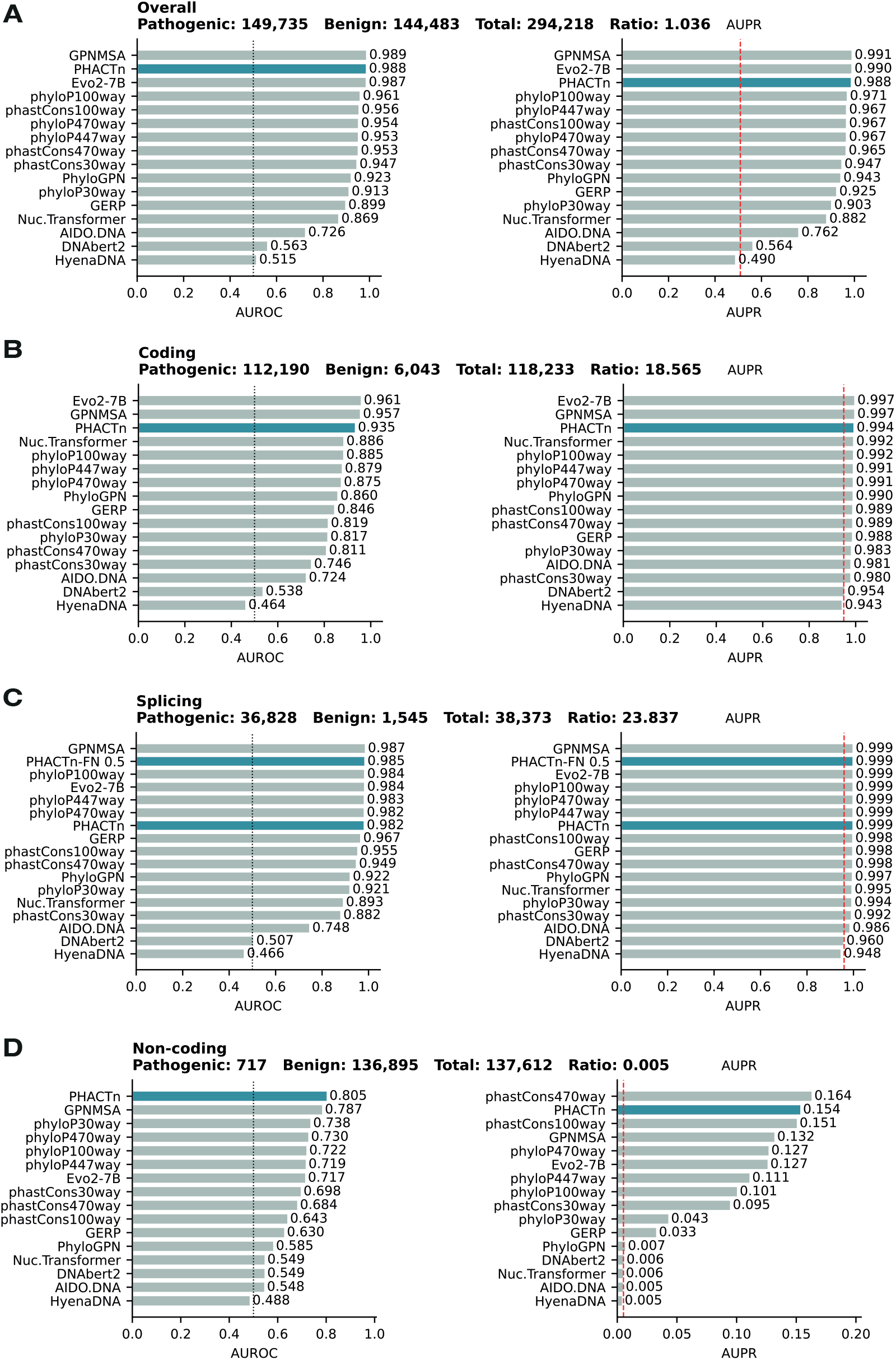
Performances of widely used benchmarking tools on the ClinVar–gnomAD variant set. AUROC and AUPR are shown for **(a)** overall, **(b)** coding (missense, synonymous, stop-gained/lost, start-lost, stop-retained), **(c)** splicing (splice acceptor/donor, donor region, 5th base, polypyrimidine tract, splice region), and **(d)** non-coding (5’/3’ UTR, intronic, intergenic, upstream/downstream, ncRNA exon, mature miRNA) variants.

**Table 1:**
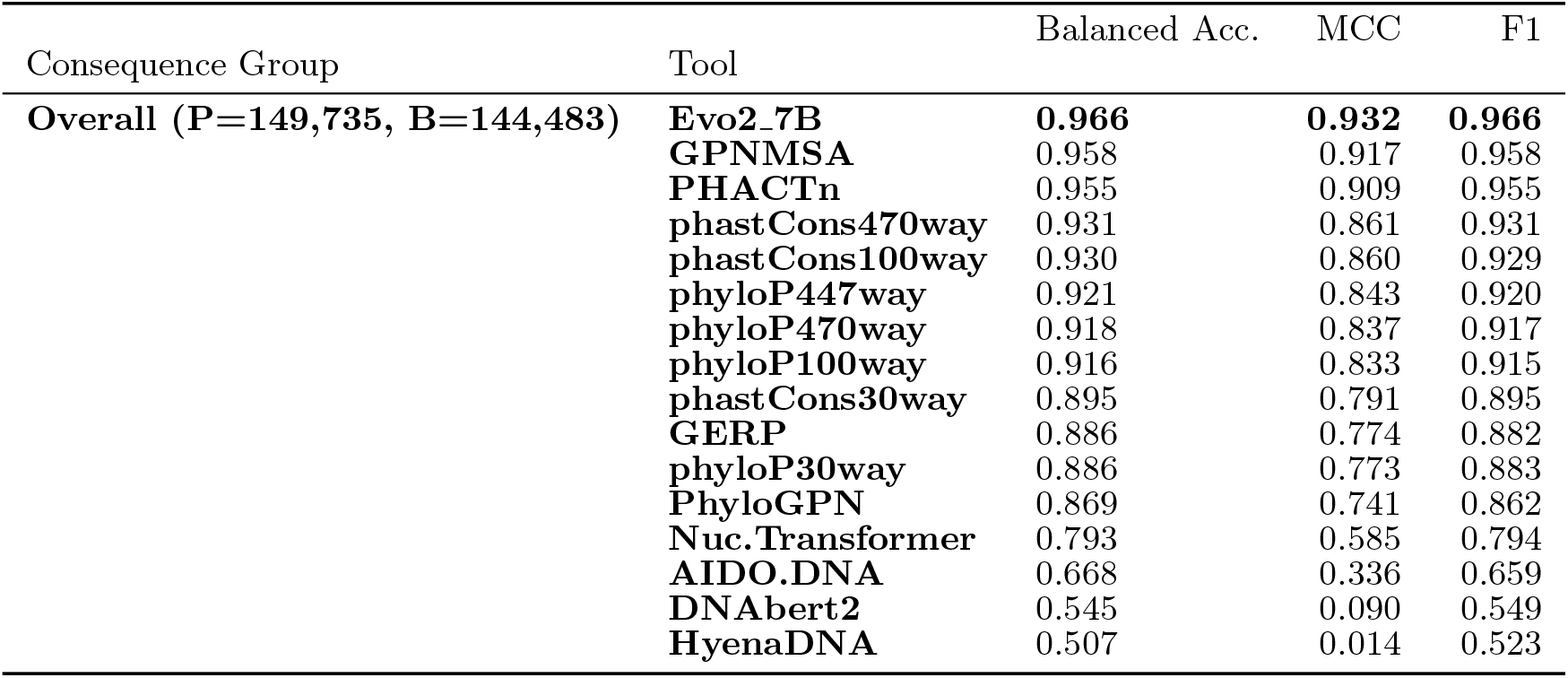
Threshold based metrics on the CG - Overall set.

On the overall CG dataset (149,735 pathogenic and 144,483 benign variants), PHACTn achieved the second-highest AUROC score among all evaluated methods, reaching 0.988, just below GPN-MSA (0.989) and surpassing Evo2-7B (0.987). A consistent pattern was observed in AUPR, where PHACTn (0.988) ranked third below GPN-MSA (0.991) and Evo2-7B (0.990), with differences of only 0.002–0.003 from the top-performing methods. For threshold-dependent metrics, Evo2-7B ranked first with F1=0.966, balanced accuracy=0.966, and MCC=0.932.

GPN-MSA followed closely with F1=0.958, balanced accuracy=0.958, and MCC=0.917, while PHACTn ranked third with F1=0.955, balanced accuracy=0.955, and MCC=0.909; differing from GPN-MSA by only 0.003–0.008 across all three metrics.

PHACTn outperformed classical conservation scores, including phyloP100way, phast- Cons100way, phyloP470way, and GERP, as well as PhyloGPN. Moreover, it is noteworthy that the HyenaDNA, AIDO.DNA and DNABERT-2 models exhibit significantly lower performance across all metrics. This suggests that, despite being large-scale DNA language models,these models fall short compared to evolutionary alignment-based methods when applied in a zero-shot manner to variant effect prediction, particularly in capturing nucleotide variability at the single-position level.

To compare performance based on variant impact, the variants in the CG dataset were annotated using Ensembl VEP [47] with the --most severe flag and categorized into three main consequence categories to achieve an adequate pathogenic-benign ratio (Table 2). The performance across the three groups is shown in Figure 3b–3d. For coding (pathogenic:benign ratio 18.6:1) and splicing (23.8:1) variant categories, we focus on AUROC, as AUPRC is inflated by the high pathogenic prevalence and is therefore less informative under these class distributions. For the non-coding category (ratio 0.005:1), where pathogenic variants constitute only 0.5% of the dataset, we report both metrics, as they answer complementary questions: AUROC reflects overall ranking ability across the full score distribution, while AUPRC more faithfully captures the practical difficulty of retrieving true pathogenic variants against a large benign background.

**Table 2:**
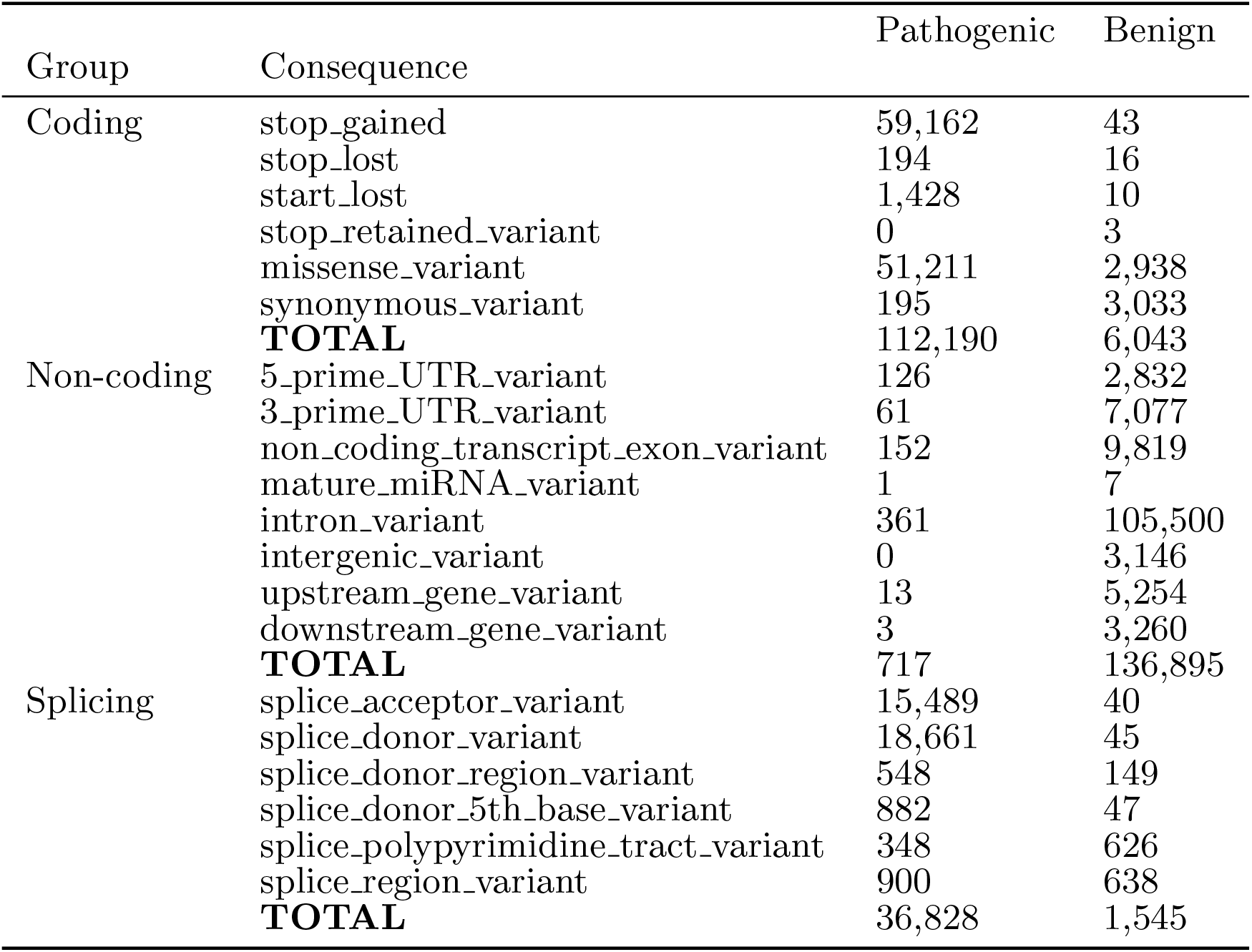
Per-consequence variant counts for Coding, Non-coding, and Splicing groups on the CG set.

**Table 3:**
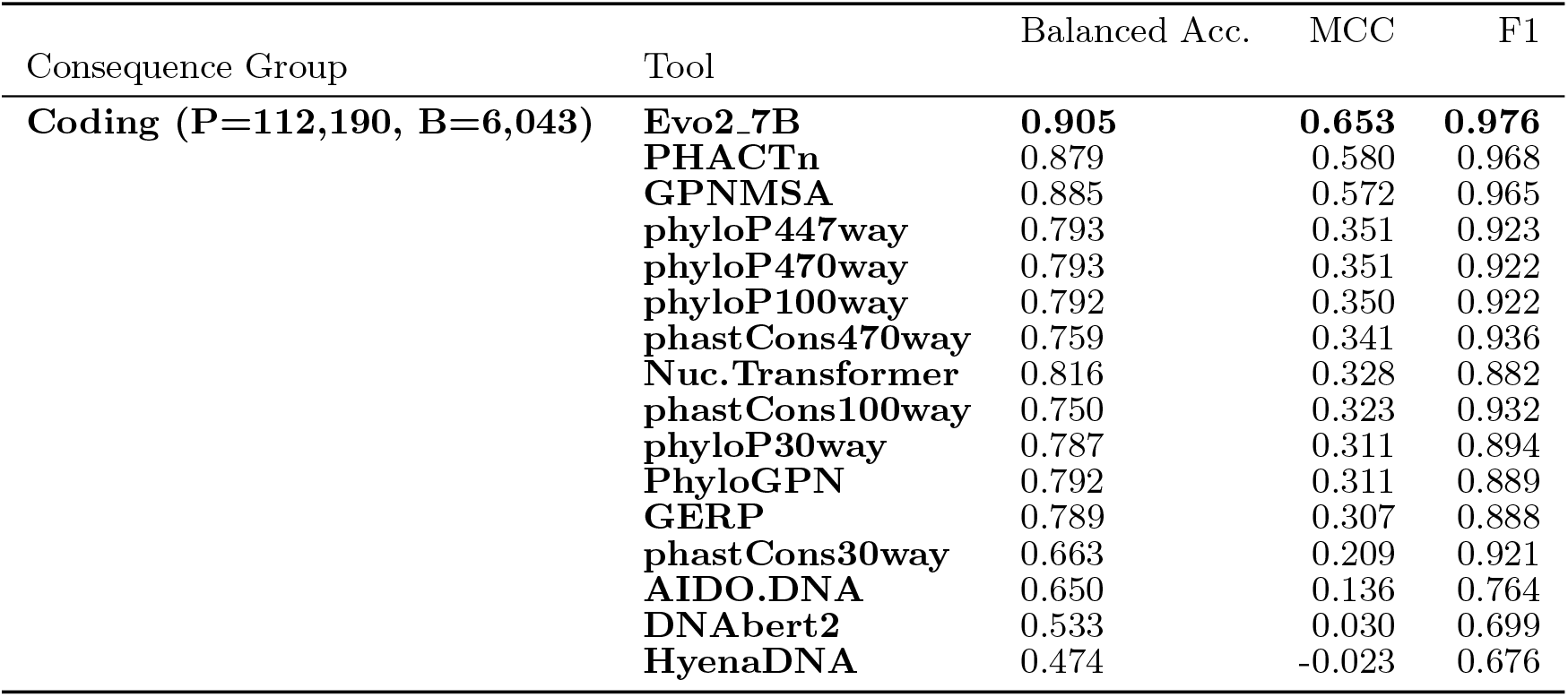
Threshold based metrics on the CG - Coding set.

In the coding group, which includes stop-gained, stop-lost, start-lost, stop-retained, missense, and synonymous variants, Evo2-7B and GPN-MSA lead with AUROC values of 0.961 and 0.957, respectively. PHACTn follows closely with AUROC 0.935. Nucleotide Transformer (0.886) and PhyloGPN (0.860) rank next, while conservation scores such as phastCons, phyloP, and GERP are solid but clearly behind (AUROC 0.811–0.885).

This performance gap may reflect fundamental differences in model architecture. Evo2 and GPN-MSA interpret sequences as contextual windows, inherently acquiring knowledge of codon structure, such as triplet patterns, reading-frame dependencies, and relationships between amino acid changes, which provides a natural advantage in coding variant prediction. However, whether gLMs such as Evo2 learn biologically meaningful relationships or simply exploit statistical regularities in massive training datasets remains debated [48]. In the case of GPN-MSA, an additional factor likely contributes to its strong performance on coding variants. GPN-MSA is trained predominantly on the top 5% most conserved genomic windows, with only 0.1% of remaining windows sampled to prevent extreme distribution shift [33]. This means the model is exposed overwhelmingly to constrained sequence contexts, including exons, splice sites, and conserved regulatory elements, while neutral and weakly functional regions are severely underrepresented. As a result, the model may develop an implicit bias toward recognizing constraint-associated sequence motifs. However, this choice may limit the tool’s generalization to genuinely novel functional signals in less conserved regions.

PHACTn, by contrast, derives its scores entirely from an explicit probabilistic model of nucleotide substitution histories across the phylogenetic tree, making its predictions directly interpretable in evolutionary terms without reliance on learned representations. Despite the architectural disadvantage of scoring each position independently, it remains competitive with others in the coding variant subset.

In the splicing category (Figure 3c), GPN-MSA leads with AUROC 0.987, followed by PHACTn-FN 0.5 with AUROC 0.985, another version of PHACTn using a distance-only Gaussian weighting with a fixed bandwidth of 0.5, rather than the default Dual-metric_4_ scheme. phyloP100way and Evo2-7B follow closely at 0.984, while the standard PHACTn (Dual-metric_4_) achieves 0.982. There is a marginal improvement of PHACTn-FN 0.5 over PHACTn (0.003 AUROC difference).

Non-coding variants represent the most challenging category for variant effect prediction, with only 717 pathogenic variants among 136,895 benign ones (Figure 3d). In terms of AUROC, PHACTn achieves the highest score of 0.805, followed by GPN-MSA (0.787). However, when evaluated by AUPR, a metric more sensitive to performance under severe class imbalance, phast- Cons470way achieves the highest score (0.164), slightly ahead of PHACTn (0.154). This likely reflects a fundamental difference between the two scores. While PHACTn evaluates each genomic position independently, without incorporating information from neighboring positions, phastCons estimates the probability that a region is conserved using Hidden Markov Models (HMMs) [31]. This method might be effective at flagging the pathogenic outliers, such as variants that fall within deeply conserved elements. Additionally, when threshold-based metrics were compared (see Table 5), it was observed that PHACTn shows improved performance over other tools in MCC and balanced accuracy. Altogether, these results suggest that, for non-coding variants, phylogenetic modeling captures constraint signals better than large-scale learned representations evaluated in this benchmark set.

**Table 4:**
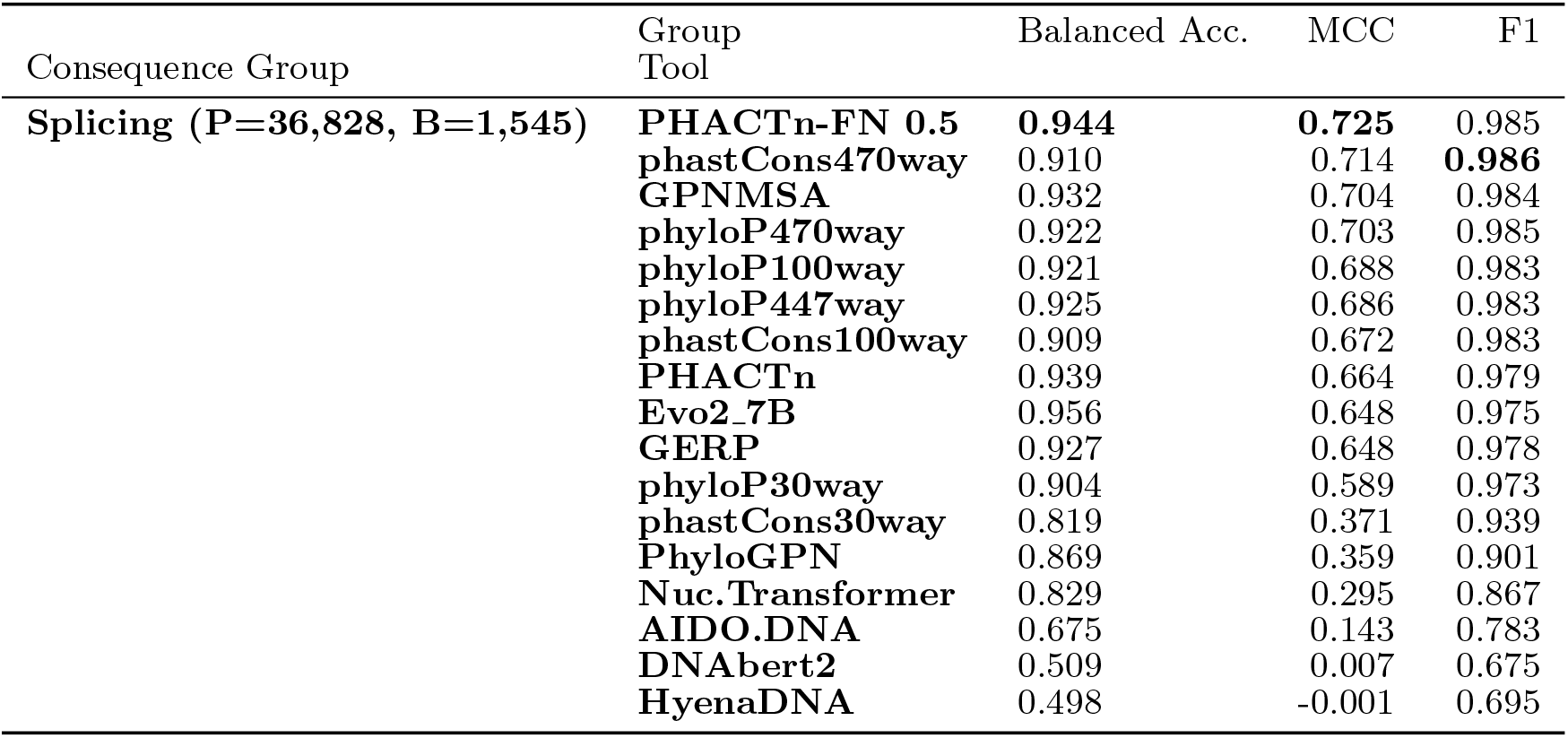
Threshold based metrics on the CG - Splicing set.

**Table 5:**
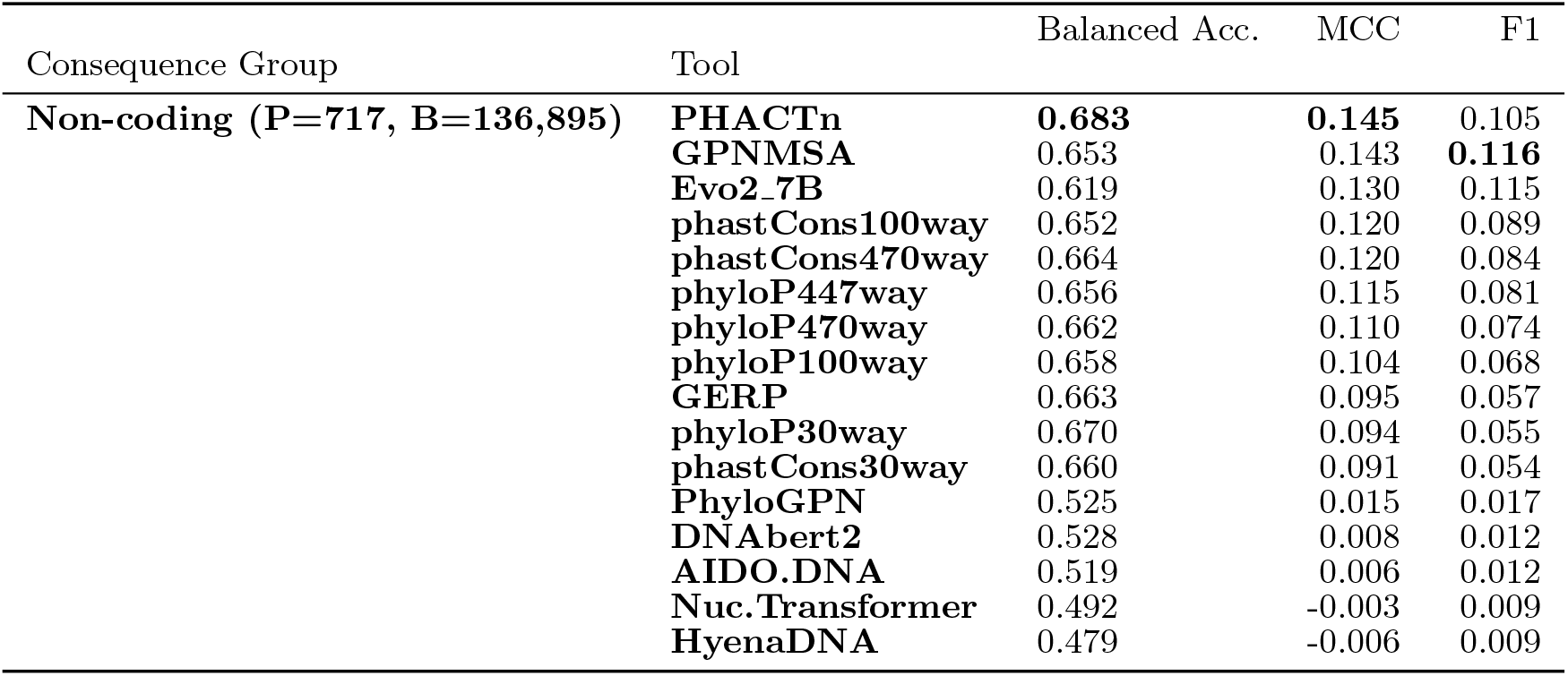
Threshold based metrics on the CG - Noncoding set.

#### 2.2.3 Evaluation on Clinically Ambiguous Variants

The variants where existing tools agree generally involve cases that carry a strong signal of evolutionary conservation, thus they are easy to classify. However, sometimes in clinically ambiguous cases, many prediction tools disagree on their effects, leading to discrepancies. Demonstrating good performance in such challenging cases, where existing methods fall short, is the strongest evidence that a method provides an independent and complementary signal that other tools fail to capture. For that reason, we created a hard-case dataset to evaluate PHACTn performance.

For the construction, phyloP and phastCons scores were each represented by a single version to avoid overrepresentation of conservation-based signals in the disagreement criterion. Specifically, the 100-way vertebrate alignment was selected as the representative, as it is derived independently from the 470-way mammalian alignment used by PHACTn, thereby preventing circularity in the evaluation. Furthermore, PhyloP scores were preferred over phastCons because they operate at single-nucleotide resolution.

PhyloP-V, Evo2, GPN-MSA, PhyloGPN, GERP, AIDO.DNA and Nucleotide Transformer scores were used in this analysis. We excluded DNABERT-2 and HyenaDNA due to their poor performance, which can be attributed to their underlying modeling limitations. DNABERT-2 relies on Byte Pair Encoding (BPE) tokenization, which compresses sequences but sacrifices the fine-grained positional detail required for single-nucleotide variant prediction [26, 49]. Furthermore, while HyenaDNA operates at a single-nucleotide resolution, it is pre-trained only on a single human reference genome [50], preventing it from learning deep evolutionary constraints.

After applying tool-specific thresholds calibrated to balance sensitivity and specificity, variants where three tools diverged from the majority prediction were designated as hard cases, accounting for 9% of the total dataset. To ensure an unbiased evaluation, PHACTn was removed from consensus voting, thus allowing an independent assessment of PHACTn’s performance on truly ambiguous variants. The results can be seen in Figure 4.

**Fig. 4:**
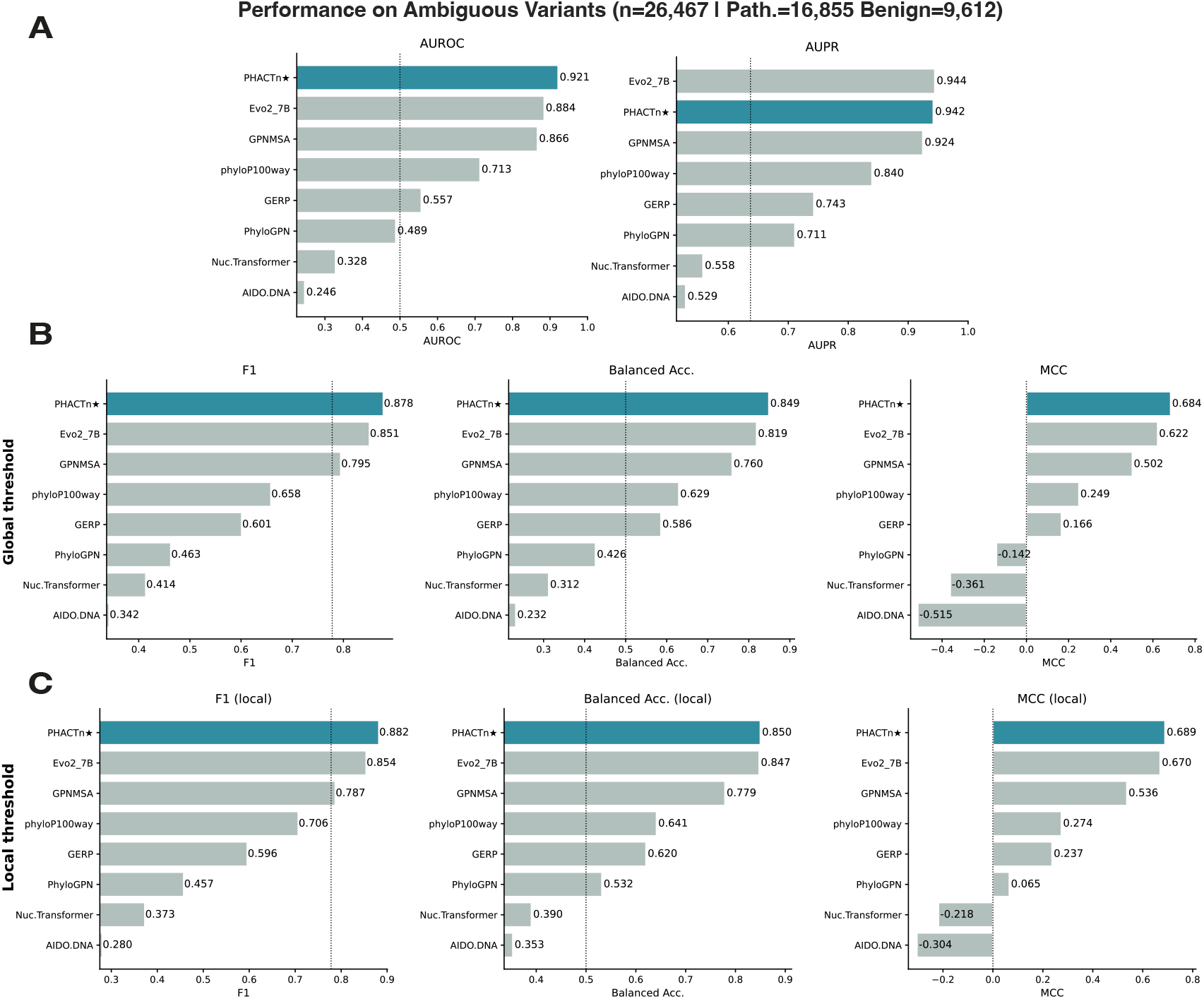
AUROC and AUPR values (a), F1 Score, Balanced Accuracy, and MCC values with thresholds calculated from the overall CG set (b) and calculated locally on the hard subset (c), for 7 benchmark tools and PHACTn on a clinically ambiguous hard-case variant set derived from ClinVar-gnomAD.

PHACTn achieved the highest AUROC (0.921), surpassing Evo2-7B (0.884) and GPN-MSA (0.866), while in AUPR, Evo2-7B led marginally (0.944) with PHACTn close behind (0.942). PHACTn’s advantage extended to threshold-dependent metrics, indicating superior calibration at the decision boundary and more accurate binary classifications. These metrics were assessed under two strategies: a global threshold already calculated from the full dataset (Figure 4b), and a local threshold calculated by maximizing the harmonic mean of sensitivity and specificity on the hard-case subset itself (Figure 4c). Under both settings, PHACTn ranked first in F1, balanced accuracy, and MCC, achieving F1=0.882, balanced accuracy=0.850, and MCC=0.689 under the local threshold. Evo2-7B performed strongly as a second across all metrics, while GPN-MSA ranked third. Conservation-based tools and sequence models such as Nucleotide Transformer and AIDO.DNA struggled considerably on this subset, with several yielding negative MCC values, indicating near-random or inverse classification on these particularly ambiguous variants. Overall, PHACTn demonstrates a consistent advantage on hard cases across both threshold-free and threshold-dependent evaluations.

#### 2.2.4 PHACTn is the Top Pathogenicity Predictor of the Mendelian Non-coding Variants

In addition to the CG variant set created for performance comparison, the TraitGym Mendelian benchmark dataset [29] was also used. In this dataset, the authors curated non-coding variants associated with 113 Mendelian diseases from OMIM by matching each positive variant with 9 negative control variants on the chromosome, consequence, and transcription start site distance. Accurate classification of non-coding variants is critical because variants in these regions can disrupt regulatory mechanisms, including transcription factor binding sites, splice donor and acceptor sequences, enhancer and silencer elements, and untranslated region-mediated post-transcriptional regulation. All of them can lead to several diseases through altered gene dosage or aberrant transcript processing.

On this set, PHACTn achieved the highest AUROC (0.914) and AUPR (0.736), followed closely by GPN-MSA (AUROC: 0.905; AUPR: 0.688) (Figure 5a). However, Evo2-7B, which performed strongly on the CG coding variant set, ranked considerably lower on this (AUROC: 0.699; AUPR: 0.332), suggesting that this model’s performance may not generalize well to non-coding variants once again.

**Fig. 5:**
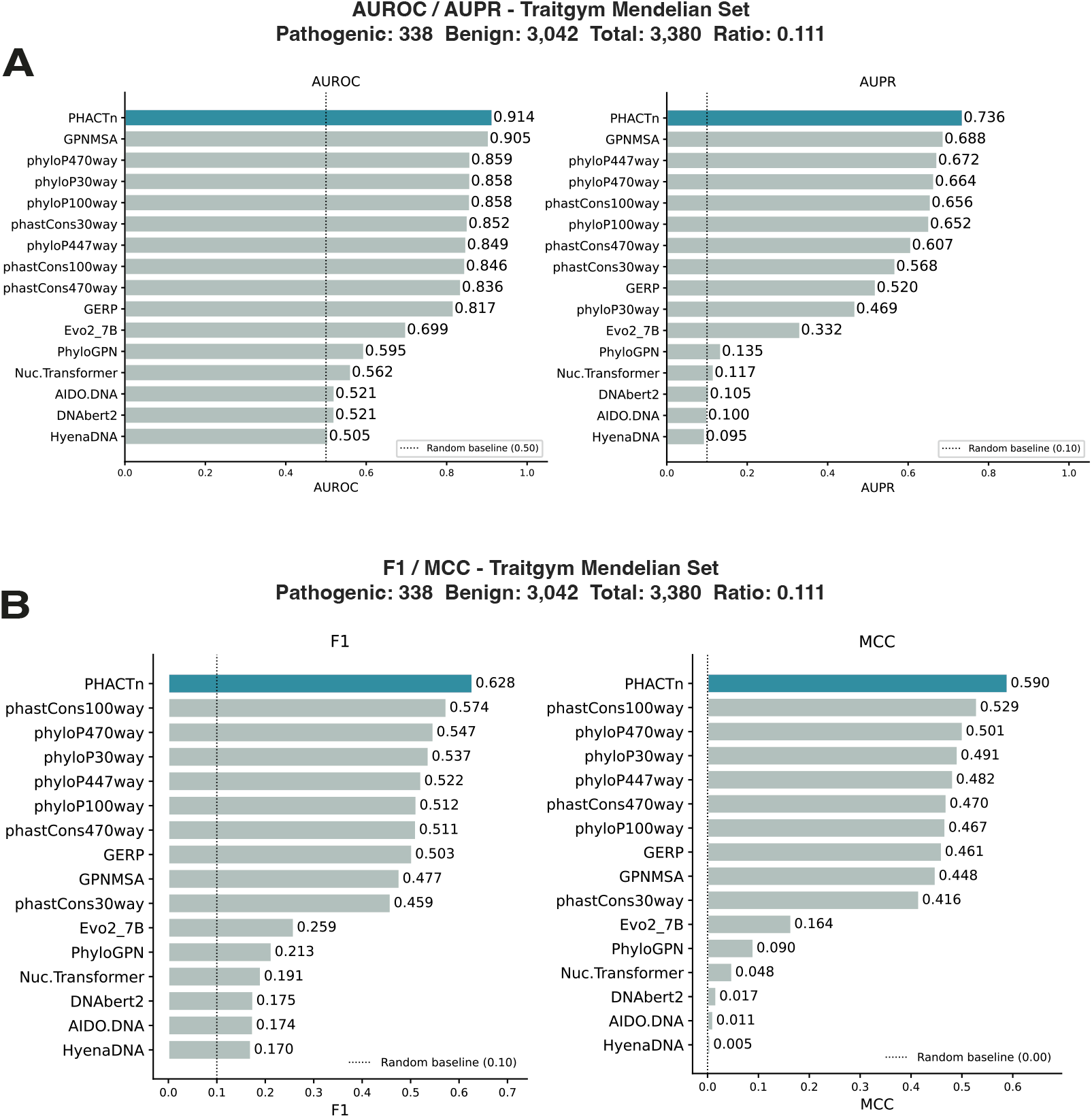
AUROC and AUPR values (a), and F1 Score and MCC values (b), of PHACTn and benchmark tools evaluated on the TraitGym Mendelian non-coding variant set (338 pathogenic, 3,042 benign; 1:9 positive-to-negative ratio).

At the best classification threshold, PHACTn again outperformed all tools in both F1 (0.628) and MCC (0.590) (Figure 5b). Among benchmark tools, conservation-based methods such as phastCons100way (F1: 0.574; MCC: 0.529) and phyloP470way (F1: 0.547; MCC: 0.501) were the strongest competitors. It may suggest that evolutionary conservation signals are particularly informative for the tolerability of non-coding variants. In contrast, large language model-based approaches showed notably weaker performance on this variant class: while GPN-MSA remained competitive in AUPRC (0.688), it fell considerably behind in threshold-based metrics, and Evo2-7B, PhyloGPN, Nucleotide Transformer, DNABERT-2, AIDO.DNA, and Hye- naDNA underperformed across all evaluated metrics. Collectively, these results demonstrate that PHACTn maintains consistent superiority across both ranking-based and threshold-based metrics even in a non-coding Mendelian disease context.

### 2.3 PHACTn Correctly Classifies Variants That State-of-the-Art Tools Fail to Predict

To identify variants that PHACTn correctly classifies while other tools fall short, seven illustrative examples were drawn from the CG dataset: the top three (Figure 6a–6c) are benign variants with high allele frequency in gnomAD, and the bottom four (Figure 6e–6h) are from ClinVar. For pathogenic variants, GPN-MSA and Evo2-7B were chosen as reference tools, as both represent the strongest competing methods on the CG benchmark. For benign variants, GPN-MSA and phyloP were used instead. By adding phyloP, we aimed to highlight the fundamental difference in how PHACTn models evolutionary constraint compared with classical position-level conservation scores. Binary labels were then assigned independently for each tool, using its optimal threshold selected to minimize the divergence between sensitivity and specificity.

**Fig. 6:**
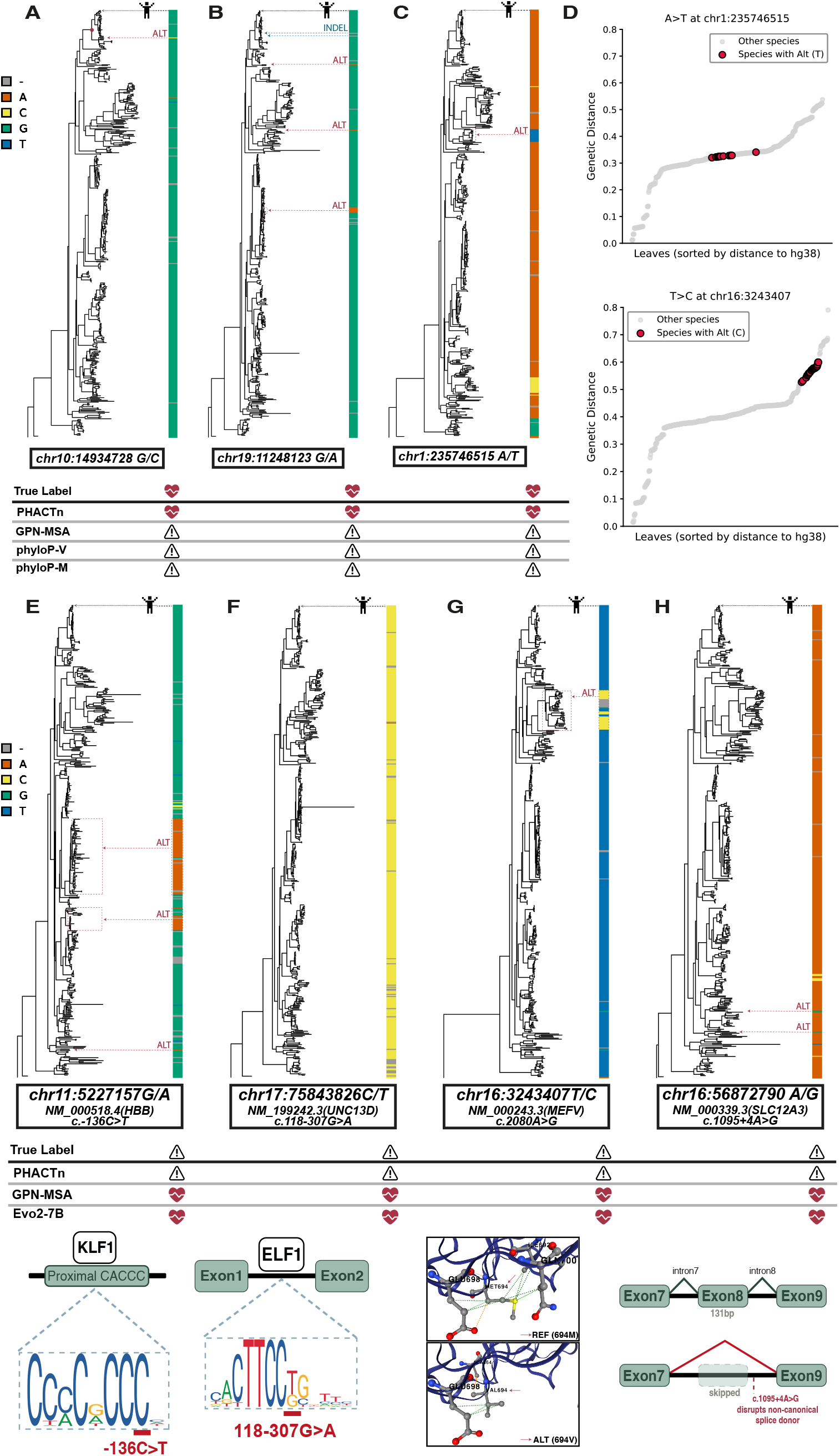
Seven variants correctly classified by PHACTn but misclassified by state-of-the-art tools. Heart and exclamation point in a triangle indicate benign and pathogenic predictions, respectively. **(a, b, c)** Three benign variants (chr10:14934728 G/C, chr19:11248123 G/A, chr1:235746515 A/T (coordinates are on positive strand)) where PHACTn correctly assigns benign labels while alignment based tools GPN-MSA, phyloP-V, and phyloP-M predict pathogenicity. **(d)** A plot showing the evolutionary distance of the species observed in the phylogenetic tree of the two examples. Species with alternative nucleotides are marked in red. The top example, chr1:235746515 A-to-T, is a benign variant with a high allele frequency on the gnomAD dataset. The true label of the botom example, chr16:3243407 T-to-C, is pathogenic. Even though their MSA patterns appear similar, the species where the alternative allele is present are closer to humans in the top example. Therefore, PHACTn considers it more tolerable and matches its label. **(e)** A pathogenic variant where PHACTn correctly predicts pathogenicity while GPN-MSA and Evo2-7B fail: NM 000518.4(HBB):c.-136C*>*T (chr11:5227157 G*>*A). KLF1 binding motif at the HBB proximal CACCC box, with the c.-136C*>*T variant disrupting a high-information-content position. **(f)** A pathogenic variant where PHACTn correctly predicts pathogenicity while GPN-MSA and Evo2-7B fail: NM 199242.3(UNC13D):c.118-307G*>*A (chr17:75843826 C*>*T). ELF1 binding motif at the UNC13D intronic regulatory region disrupted by c.118-307G*>*A. **(g)** A pathogenic variant where PHACTn correctly predicts pathogenicity while GPN-MSA and Evo2-7B fail: NM 000243.3(MEFV):c.2080A*>*G (chr16:3243407 T*>*C). DynaMut2-predicted structural consequence of MEFV p.Met694Val (c.2080A*>*G) on the AlphaFold model, showing altered residue interactions in the B30.2 domain. **(h)** A pathogenic variant where PHACTn correctly predicts pathogenicity while GPN-MSA and Evo2-7B fail: NM 000339.3(SLC12A3):c.1095+4A*>*G (chr16:56872790 A*>*G). Splicing diagram of SLC12A3 illustrating exon 8 skipping caused by disruption of the non-canonical splice donor by c.1095+4A*>*G.

The first example in Figure 6a illustrates a tolerable variant that is incorrectly predicted as conserved by phyloP-V (100 vertebrate alignment), phyloP-M (470 mammalian alignment), and GPN-MSA. A single G-to-C substitution is observed at a conserved position, which alone would typically not be sufficient to indicate tolerability. However, the phylogenetic tree reveals that this substitution occurred in a lineage closely related to humans, which PHACTn appropriately captures when assigning its tolerability score.

The second example in Figure 6b demonstrates a similar case, but with the additional contribution of indels. Three independent G-to-A substitutions are observed at this position; however, unlike the previous example, these occur in lineages more distant from humans and thus contribute only modestly to the final score. More importantly, two indels are present within the same clade as humans, and these increase the tolerability score of the four nucleotides after the weighted gap frequency step.

The third example in Figure 6c appears ambiguous at first glance, given a single clade-specific substitution occurring at a relatively large phylogenetic distance from humans. While superficially similar to the pathogenic example below (Figure 6g), the key distinction lies in the phylogenetic proximity of the substitution node to humans, which is closer than its pathogenic counterpart (Figure 6d). This difference is sufficient for PHACTn to correctly assign a benign label to chr1:235746515 (A-to-T), while accurately classifying the lower variant as pathogenic, underscoring the importance of precise phylogenetic weighting in distinguishing neutral from deleterious variation.

Having examined cases where PHACTn correctly identifies benign variants that conservation-based tools misclassify as deleterious, we now turn to the complementary scenario: pathogenic variants that GPN-MSA and Evo2-7B fail to detect, yet PHACTn correctly flags. One illustrative example (Figure 6e) NM 000518.4(HBB):c.-136C*>*T (G-to-A in positive strand), located in the proximal CACCC box in the promoter of the beta-globin gene [51–53]. It has been proven that mutations in this promoter area decrease beta-globin transcription [54]. The variant -136C*>*T has also been reported in a beta-thalassemia patient [55]. Among the transcription factors that regulate HBB expression, KLF1 plays the most critical role. It binds the proximal CACCC box of the *β*-globin promoter and drives high-level erythroid-specific transcription; KLF1 knockout mice fail to activate *β*-globin expression and develop a severe, lethal *β*-thalassemia-like anemia [54].

To further corroborate the functional relevance of this variant, we downloaded the KLF1 position weight matrix (MA0493.2) from the JASPAR database [56] and performed motif scanning using FIMO [57] with a 20 bp flanking sequence on each side of the variant (threshold: *p <* 10*^−^*^3^). FIMO identified a significant KLF1 motif hit spanning positions 20–28 of the query sequence (score = 13.13, *p* = 3.1 10*^−^*^5^), placing the c. 136C*>*T variant precisely at position 2 of the KLF1 binding motif, one of the highest-information-content positions in the motif logo (Figure 6e). This indicates that the variant may directly disrupt a highly conserved nucleotide within the KLF1 recognition sequence, providing a mechanistic explanation for its pathogenicity.

In the 470-mammal alignment, ’A’ nucleotides are observed across nearly 80 species, a count that might superficially suggest tolerance for the alternative allele. However, PHACTn correctly identifies this as pathogenic because these observations trace back to only 3 independent evolutionary events, as can be seen in the figure. Critically, all three events occur far from the human lineage. Under PHACTn’s sharply decreasing distance weight, the signal is dominated by the dense conservation of ’C’s among species closely related to humans. This example highlights an important limitation of approaches that count raw nucleotide frequencies across species without accounting for phylogenetic structure. A näıve count of 80 ”A” observations could easily mislead a model into treating the position as tolerant, whereas PHACTn’s weighting correctly recognizes that these observations reflect a small number of ancient, distant substitution events rather than widespread functional tolerance near the human lineage.

Second example (Figure 6f), the variant NM 199242.3(UNC13D):c.118-307G*>*A (C-to-T in positive strand) is located within intron 1 of the UNC13D gene. The studies show that this variant reduces RNA expression by disrupting the binding site for ETS family transcription factors [58, 59]. In general, mutations in UNC13D, which encodes a protein essential for lytic granule fusion in NK cells and cytotoxic T lymphocytes, cause Familial Hemophagocytic Lymphohistiocytosis type 3 (FHL3), characterized by defective lymphocyte cytotoxicity [60]. By analogy with the neighboring c.118-308C*>*T variant, for which disruption of ELF1 binding and resulting loss of STAT4 and BRG1 recruitment have been experimentally demonstrated [61, 62]. Moreover, c.118- 307G*>*A is predicted to impair the same regulatory axis, though direct experimental evidence for STAT4/BRG1 disruption at this specific position has not yet been reported.

To check whether we observe conservation at the ELF1 binding site, as in the previous one, we used the same approach. We downloaded all available ELF1 position weight matrices from JASPAR [56], ran FIMO [57] with 20bp flanking sites, and identified the highest-scoring motif (MA0473.1, score = 12.3673, *p* = 3.38 10*^−^*^5^). The highlighted position in the motif logo (Figure 6f) reveals that the ELF1 motif strongly favors C and A on the positive strand, corresponding to G and T on the negative strand. The c.118-307G*>*A substitution introduces an adenine on the negative strand, a nucleotide that is essentially absent from the position weight matrix, indicating severe intolerance at this position. This finding is consistent with PHACTn correctly flagging the variant as pathogenic: PHACTn observes no thymine at this position across mammals, providing a strong phylogenetic signal of intolerance. Also, the indels don’t provide strong signal toward tolerability because its weighted contribution is low, given most indels are shown in species far from human. Furthermore, in the broader 100-vertebrate whole-genome alignment, thymine appears in only two species, both of which are birds, a distantly related taxon to humans, strengthening the idea that the alternate allele is not tolerated.

Another case (Figure 6g), the variant NM 000243.3(MEFV):c.2080A*>*G (p.Met694Val, T-to-C in positive strand), causes Familial Mediterranean Fever (FMF) through gain-of-function and is in fact the most common pathogenic variant for this disease [63]. This variant is in the B30.2 domain of the MEFV gene, which encodes the pyrin protein. Chae et al. showed that the corresponding region forms distinct loops that create an interface that interacts with caspase-1, as demonstrated by computational docking, and claimed that M694V physically disrupts this interaction, reducing pyrin’s ability to bind and inhibit caspase-1, thereby releasing the inhibitory brake and enabling uncontrolled IL-1*β* production [64]. In one study, hydrophobic clusters were identified within this domain that may function as a binding site [65]. In a more recent study, M694V was shown to behave as if the B30.2 domain is deleted, providing evidence for caspase-1-independent regulation of pyrin [66].

To assess whether M694V affects pyrin stability, we used DynaMut2 [67] on the AlphaFold structure of the protein [68] (id: AF-O15553-F1-v6). The predicted stability change was 1.68 kcal/mol, indicating a destabilizing effect. The interactions of the reference and alternate alleles within the protein structure can be seen in Figure 6g, zoomed to the position of interest.

Even though this variant is well-defined clinically, Evo-2 and GPN-MSA fail to predict its pathogenicity, whereas PHACTn correctly classifies it as pathogenic through a more refined interpretation of the mammalian alignment. Indeed, the alternative nucleotide G is observed, yet PHACTn correctly identifies the variant as pathogenic because G is the consensus nucleotide within that clade and represents lineage-specific fixation rather than functional equivalence with the human context.

A fourth example (Figure 6h) extends PHACTn’s advantage to the category of splice-site variants. The variant NM 000339.3(SLC12A3):c.1095+4A*>*G, classified as likely pathogenic in ClinVar, causes Gitelman syndrome by disrupting the non-canonical donor splice site of intron 8, leading to exon 8 skipping by silencing the 131 base pairs and leading to frameshift, as confirmed by an in vitro midigene assay (Figure 6h) [69, 70]. Notably, SpliceAI and Pangolin assign negligible scores at this position [71] (Acceptor Loss: 0.05, Donor Loss: 0.01, Acceptor Gain: 0.01, Donor Gain: 0.01, Splice Loss: 0.07, Splice Gain: 0.01), missing the experimentally validated splicing disruption. In the 100-vertebrate alignment used by GPN-MSA, the alternative allele G is observed in 17 species distributed across three phylogenetically distant groups: ray-finned fishes (n=1), lobe-finned vertebrates (n=5), and birds (n=11). These represent at most 3 independent evolutionary events, both occurring in lineages far from mammals. PHACTn, operating on 470 mammals, finds G essentially absent across the entire mammalian clade, the position is nearly invariant among mammals, and the two independent appearances of G occur exclusively in distant lineages. Under PHACTn’s harshly decreasing distance weight, these distant observations contribute minimally to the score, and the conservation of A among close mammalian relatives dominates the pathogenicity signal.

Collectively, these seven examples illustrate the core advantage of PHACTn’s phylogenetic weighting strategy. Rather than treating all observed substitutions equally across the alignment, PHACTn explicitly accounts for evolutionary distance and distinguishes lineage-specific fixation from true functional equivalence, ensuring that rare, distant events or clade-specific substitutions do not obscure the strong conservation signal present among species closely related to humans.

### 2.4 PHACTn Achieves Competitive Performance with Minimal Complexity

Most benchmarks evaluate variant effect predictors solely on predictive accuracy, leaving aside an important question: can the tool realistically be used by the broader research community? Large foundation models may achieve strong performance, but they come with practical barriers because they require expensive GPUs to run inference, consume months of training time on massive datasets, and produce scores that cannot be interpreted or traced back to biological principles. For most research groups without access to large-scale compute infrastructure, these tools are effectively out of reach. A truly useful variant effect predictor must not only perform well, but must also be accessible, efficient, and transparent in how it arrives at its predictions. On the other hand, PHACTn is accessible, efficient and transparent.

In terms of parameterization, compared to the hundreds of millions or billions of parameters in contemporary models, PHACTn uses just 4 interpretable parameters: the weighting function, the normalization strategy, the gap correction threshold t, and the slope s. This difference spans more than nine orders of magnitude (Figure S4b).

Beyond parameter count, the practical barriers imposed by large models are substantial. Evo2 requires high-end GPU infrastructure, such as NVIDIA A100 or H100 accelerators, to run inference at a genome scale, and training such models requires months of compute time on large GPU clusters. By contrast, PHACTn requires no GPU, no training phase, and no specialized hardware; it runs on standard CPUs using alignments and a phylogenetic tree.

Beyond scale, PHACTn’s evolutionary scope is narrower than that of its competitors (Figure S4b). GPN-MSA is trained on a 100-vertebrate whole-genome alignment spanning the full vertebrate clade, and Evo2 was pre-trained on sequences drawn from all domains of life. PHACTn, by contrast, operates exclusively on the 470-way mammalian alignment. With that, PHACTn achieves competitive or superior performance by taking advantage of a more limited but higher-quality evolutionary signal rather than compensating for alignment noise with learned parameters.

Crucially, PHACTn’s scores are fully interpretable. For any given variant, the contribution of each species and each substitution event to the final tolerance score can be traced directly through the phylogenetic tree traversal. This transparency stands in sharp contrast to the black-box representations produced by transformer-based and autoregressive sequence models, whose internal reasoning cannot be inspected or audited.

Despite these constraints, no training, no learned parameters, no GPU, mammal-only alignment, PHACTn remains competitive with or superior to GPN-MSA and Evo2-7B across overall, splicing, and non-coding variant categories, and ranks first in AUROC, F1 and MCC on the hard-case clinically ambiguous subset.

### 2.5 PHACTn Reaches Full Potential After Excluding Alignment-Blind Variants

A limitation shared by all alignment-based variant effect prediction methods is their dependence on the information available within the multiple sequence alignment. At positions where the alternative allele is entirely absent across the 470-mammal whole-genome alignment with few gaps, PHACTn assigns low tolerance scores, the correct evolutionary interpretation when the allele has never been observed in any lineage. However, we identified a subset of such variants that, despite being absent from the alignment, are common in the human population (gnomAD AF *>* 0.01), creating a direct conflict between the evolutionary signal and observed population frequency. Since PHACTn only sees the evolutionary signal, it systematically misclassifies these as pathogenic.

Several biological explanations may account for these cases. A variant that appears conserved across species and is predicted to be pathogenic may persist in the population due to co-evolution with a compensatory or suppressor mutation that restores fitness and buffers its functional impact [72–74]. Alternatively, such alleles may have risen to high frequency through positive selection specific to the human lineage, local adaptation, genetic drift, or demographic expansion [75]. Human-specific duplications or structural rearrangements may also alter the functional landscape in which a variant operates, leaving cross-species conservation comparisons less informative [76, 77]. Finally, technical artifacts, including poor whole-genome alignment in repetitive and low-complexity regions, as well as incomplete reference genomes of the non-human species, may further confound the signal. All of these scenarios represent fundamental limitations of any phylogeny-based approach that assumes cross-species conservation directly translates to functional constraint within a single lineage.

We identified and removed these problematic cases (alternative frequency = 0, gap frequency ≤ 0.3, and benign; total 2128 variants) to represent an informative and fair category of variants. The removal criterion is based purely on alignment properties and population genetics; PHACTn’s output played no role in identifying which variants to remove. Moreover, GPN-MSA and phyloP/phastCons exhibit the same blind spot at these positions, and removing them does not selectively benefit PHACTn. Since the full benchmark results, including these variants, are already reported above, this analysis provides a complementary view of PHACTn’s performance on evolutionarily tractable variants rather than a replacement for the primary evaluation.

The results are presented in Figure S5. After excluding these conflicting benign variants, PHACTn achieves the top performance across all evaluated metrics except the splicing group, which is only 0.03 behind GPN-MSA. On the cleaned overall dataset (149k pathogenic and 142k benign variants), PHACTn reaches an AUROC of 0.991 and AUPR of 0.993, surpassing GPN-MSA (0.990 AUROC; 0.992 AUPR) and Evo2-7B (0.987 AUROC; 0.991 AUPR) (Figure S5a).

Collectively, these results demonstrate that PHACTn’s performance on the full benchmark is not a reflection of a weakness in phylogenetic modeling, but rather an artifact of including variants that lie outside the informative range of the tool.

## 3 Discussion

In this study, we introduced PHACTn, a phylogeny-aware probabilistic method for predicting single-nucleotide variant tolerability across the entire human genome. We tested PHACTn on a ClinVar-gnomAD variant set, its hard-case subset, and the TraitGym benchmark for non-coding variants. Overall, our analyses showed that PHACTn afforded superior predictive performance compared to well-known genomic conservation scores (PhyloP, PhastCons, GERP) and DNA language models (Nucleotide Transformer, DNABERT-2, AIDO.DNA, HyenaDNA). On the other hand, our performance was competitive against alignment-based genomic language model GPN-MSA and large-scale foundational genomic model Evo2. This is noteworthy because PHACTn is a training-free method with only four interpretable parameters that relies solely on probabilistic phylogenetic tree traversal and ancestral history, yet matched models with billions of learned parameters.

Importantly, the benchmark advantage of GPN-MSA should be interpreted with some caution. Its loss function directly incorporates phastCons and phyloP scores as position-specific weights, upweighting conserved positions and downweighting repetitive or neutral regions during training [33]. This does not constitute data leakage in the strict sense, as no variant labels were used during training, but it does complicate the interpretation of benchmark performance: it remains difficult to disentangle whether GPN-MSA’s advantage reflects genuinely learned biological representations or a well-calibrated amplification of evolutionary conservation signals already embedded in its objective function. PHACTn, by contrast, relies on an explainable method to account for why it flags a variant as tolerable or deleterious.

In non-coding variant prediction, advantage of PHACTn is obvious. On both CG dataset and the TraitGym Mendelian set, it outperformed all benchmark tools in at least two of the threshold-based metrics which are important in evaluating the performance in imbalanced data. The effects of non-coding pathogenic variants are difficult to classify because they show weak patterns unlike coding regions or splice sites and they have poorly defined landscapes. PHACTn’s success in this category suggests that the phylogeny-aware probabilistic model is able to capture the informative evolutionary signals. In coding regions, on the other hand, it performed competitively but couldn’t surpass context aware language models because they may inherently learn codon structure patterns and reading-frame dependencies. This is anticipated for a position-independent strategy and may indicate that PHACTn is most beneficial as a complementary score in coding circumstances.

The hard-case analysis further underscores PHACTn’s independent and complementary contribution to variant effect prediction. These variants are particularly challenging because existing tools capture different aspects of genomic context, such as conservation, sequence grammar, and alignment frequency. Although each signal is individually informative, they point in different directions, leading tools to disagree. PHACTn’s superiority in AUROC, F1 and MCC on this subset suggests that phylogenetic independence carries a signal orthogonal to what other approaches capture. Combining PHACTn with contextual sequence models could therefore assemble a stronger predictor, particularly for the ambiguous cases where current methods most frequently disagree.

Despite its overall strong performance, PHACTn has some limitations and failure cases. One of them concerns positions that are fully conserved across mammals yet exhibit high allele frequencies in gnomAD, representing a particular challenge: PHACTn assigns low tolerance scores by design at such sites, while population prevalence suggests these variants are likely benign. Several biological and technical mechanisms may underlie this discordance, including compensatory epistasis buffering the functional impact of an otherwise deleterious change, human lineage-specific positive selection or demographic processes elevating allele frequencies, structural genomic rearrangements altering the functional context in which a variant operates, and artifacts arising from poor cross-species whole-genome alignment or incomplete non-human reference genomes. These highlight an inherent limitation of alignment and phylogeny-based constraint scores.

Another challenging category is positions with high gap frequencies (higher than 60-70%) that PHACTn assigns highly tolerable scores, but actually annotated as pathogenic in ClinVar. This may reflect technical artifacts arising from poor alignment quality. More broadly, PHACTn’s accuracy is mainly dependent on the quality of the multiple sequence alignment; regions with poor orthology assignment, low species coverage, or alignment artifacts will inevitably give less reliable scores.

A notable limitation of PHACTn is that it does not make cell-type-specific predictions. While it achieves high accuracy across a broad range of variant categories, its tolerability score reflects millions of years of evolutionary constraint across species, producing a single context-agnostic signal. Yet the functional impact of non-coding variants is often deeply context-dependent, as a variant may disrupt an enhancer that is only active in a specific tissue, leaving its effect invisible to purely evolutionary metrics. Sequence-to-function models [78–80] take a complementary approach by learning to map DNA sequences to thousands of cell-type-specific functional read- outs, thereby predicting how a variant reshapes gene expression in a given tissue. Combining PHACTn scores with such context-aware signals is a natural next step, and hybrid approaches of this kind could offer a richer and more mechanistically grounded framework for interpreting complex-trait variants, one that unites evolutionary constraint with tissue-specific regulatory logic.

An additional limitation arises from the model that is used in the ancestral reconstruction step, which assumes equal substitution rates and equal base frequencies across all positions. This model does not account for site-specific rate variation or local nucleotide composition biases, meaning that positions embedded in GC-rich regions, repetitive elements, or regions with elevated mutation rates may receive ancestral probability estimates that do not accurately reflect their true evolutionary history. More principled substitution models that incorporate genomic context, such as position-specific rate heterogeneity, could potentially improve the accuracy of ancestral reconstruction and, consequently, the reliability of PHACTn scores in these regions.

In conclusion, PHACTn offers a principled, transparent, and computationally accessible approach to variant effect prediction that requires no training data, no learned parameters, and no gene-level annotations. Future work integrating PHACTn scores with local sequence context or functional genomics annotations may further extend its utility to complex-trait variants and coding regions, where position-level independence and alignment quality are limiting factors.

## 4 Methods

### 4.1 Generation of Genomic Input Blocks

This study utilized 470 mammalian whole-genome alignments (WGA), their phylogenetic tree, and the human reference genome (GRCh38/hg38), all retrieved from the UCSC Genome Browser (Hiller Lab Multiz 470way) [42]. Each human chromosome was processed in parallel to ensure consistency across large genomic regions.

The downloaded MAF-format alignments were first converted to FASTA using the msa view command in the PHAST toolkit [81], and then, to maintain consistency with the reference coordinate framework, the alignment columns corresponding to gaps in the human sequence were removed. Before continuing with the next step, the human sequence in the alignment was double-checked against the reference genome to prevent conflicts. Then, human sequences were scanned per chromosome, and regions containing ambiguous bases (N/n) were marked to identify regions consisting only of standard bases (A, C, G, T). To optimize calculations for the next stage, the alignment was divided into windows, and the ambiguous bases observed in humans were used to define the boundaries, with a maximum length of 10 kb and a minimum of 100 bases. In cases where the sequence was naturally shorter than this threshold, they were merged with adjacent segments while preserving the ambiguous bases between them to meet the minimum length requirement. This approach prevented the use of excessively short and poor alignments, so that each resulting segment matched with the reference genomic region with high reliability. As a result, a collection of Multiple Sequence Alignments (MSA) was saved as an independent FASTA file for efficient batch processing for downstream analyses.

### 4.2 Preprocessing and Ancestral State Reconstruction

At the start of the workflow, each input block underwent an additional preprocessing step to remove species whose sequences were composed entirely of gap characters. To ensure internal consistency, the phylogenetic trees were pruned to retain only the taxa present in the filtered alignments and were left unrooted. If an alignment has fewer than 10 species, they were removed completely.

After the tree and alignment blocks were ready, ancestral state reconstruction (ASR) was done with IQ-TREE2 [40] with the model of equal substitution rates and equal base frequencies. The alignment is additionally encoded into a binary state (gap (0) / character (1)), and the presence/absence of character probability for each position at every internal node is calculated in the second ASR step. This indel reconstruction used the phylogenetic tree that was generated from the nucleotide ASR with fixed topology and branch lengths. As a substitution model, the Jukes-Cantor 2-state (JC2) was used, which is the binary version of the nucleotide ASR model.

### 4.3 Gap-aware weighting of ancestral states

The results of both ASR steps are combined using the following formula to compute gap-aware weighted ancestral probabilities:

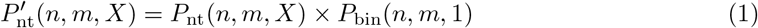

Here, *n* represents the position, *m* represents the node, and *X ∈ {A, T, G, C}* represents the nucleotide. *P_nt_*(*n, m, X*) is the probability derived from the nucleotide ASR; *P_bin_*(*n, m,* 1), on the other hand, represents the character probability generated by the binary ASR for that position and node. 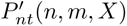, which is the product of these two probabilities, provides the weighted nucleotide probability adjusted for the gap condition.

### 4.4 PHACT - Phylogeny-Aware Computing of Tolerance Framework

The principle of PHACTn is summarized with pseudocode in Algorithm 1. As input, PHACTn takes gap-aware probability states, an alignment, and a branch-length-optimized phylogenetic tree generated from the ASR step. Starting from any selected query sequence, in this study, *Homo sapiens*, it traverses the tree, calculates and records the probability differences for each nucleotide at each connected node.

#### Algorithm 1 PHACT - Phylogeny-Aware Computing of Tolerance Framework

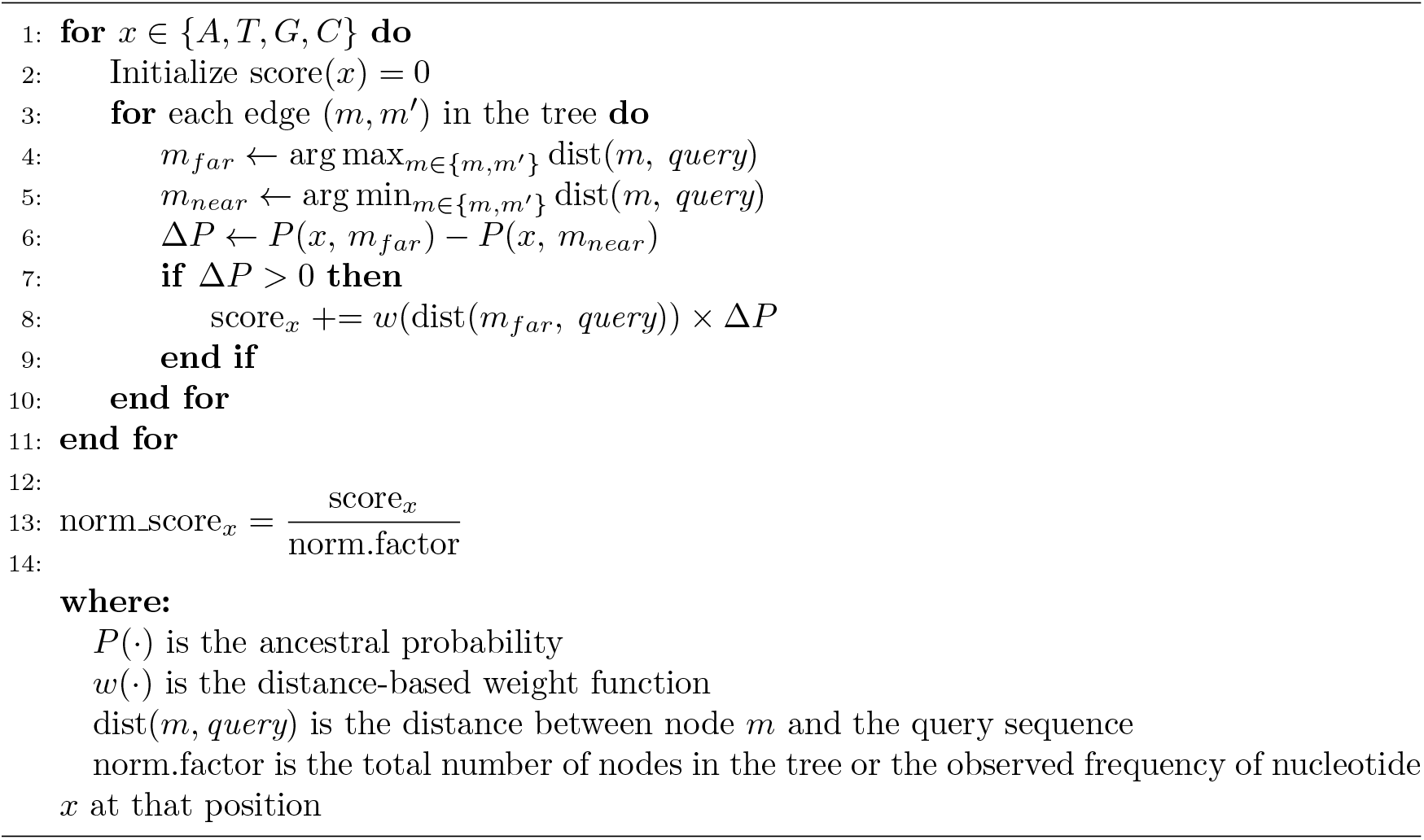

Since the probability differences accumulate during tree traversal, to avoid obtaining a larger tolerance value when a larger tree is used, the raw scores were normalized in two ways: by the total number of nodes and by the observed frequency of the query nucleotide at that position across the alignment. While the first normalization was designed solely to eliminate bias caused by tree size, the second was also attempted to provide a measure of positional diversity.

Finally, after normalizing, raw values become too small for direct biological interpretation. It is resolved by transforming the score to a continuous [0, 1] scale with Equation 2, where higher values indicate greater tolerance and lower values signal likely functional disruption.

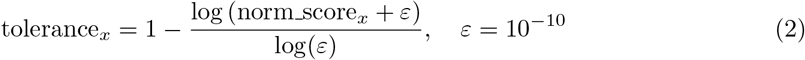

### 4.5 Weighted Gap Frequency Correction

The weighted gap frequency (WGF) is computed as the weighted average of gap indicators across all leaves of the phylogenetic tree, as shown in Equation 3:

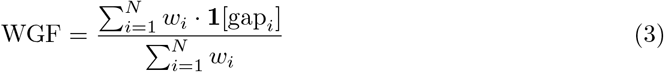

where *N* is the total number of leaves, **1**[gap*_i_*] is an indicator function that equals 1 if leaf *i* carries a gap at that position and 0 otherwise, and *w_i_* is the weight assigned to leaf *i* using the weighting function:

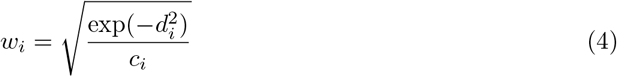

where *d_i_*is the cumulative branch length distance from leaf *i* to *Homo sapiens* and *c_i_* is the number of internal nodes on the path between them. The human leaf itself is excluded from the summation by setting *w*_human_ = 0.

Then, the gap correction term is included as follows:

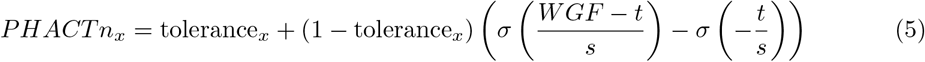

where,

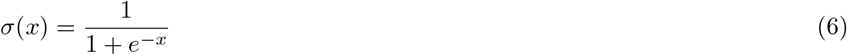

where *t* is the activation threshold, *s* controls the sharpness of the transition, and *WGF* is the weighted gap frequency defined in Equation 3. The sigmoid term in the formula ((*WGF t*)*/s*) acts as a soft activation function that determines the strength of the correction based on the observed WGF at that position. When WGF remains below the threshold value *t*, the sigmoid produces a small value, and the correction remains weak; when *WGF* exceeds *t*, the correction gradually strengthens. To ensure that the correction is exactly zero when no gap evidence is present, the baseline value of *sigmoid*(*t/s*) corresponding to *WGF* = 0 is subtracted. This process prevents an artificial upward shift in scores, thereby avoiding a systematic bias toward the benign direction.

Finally, the correction is scaled by the (1 *tolerance_x_*) factor. This ensures that the magnitude of it is inversely proportional to the current score: positions with scores already close to 1 receive little or no correction, while low-scoring positions, where gap evidence is most informative, receive the largest correction. The adjusted score remains within the [0, 1] range.

The threshold parameter *t* was set to 0.30 based on the observed distribution of WGF values across the curated ClinVar-gnomAD variant set (see 4.7). Pathogenic variants exhibited consistently low WGF values (median = 0.014, 75th percentile = 0.024), indicating that truly constrained positions are rarely gapped across the mammalian phylogeny. In contrast, benign variants showed substantially higher WGF values (median = 0.424, 25th percentile = 0.158), reflecting evolutionary relaxation. A threshold of *t* = 0.30 thus lies in the natural separation between the two distributions. The slope parameter *s* = 0.05 was selected to produce a sharp but continuous transition around this threshold, preventing abrupt discontinuities in the score while maintaining sensitivity to positions where gap frequency exceeds the threshold.

### 4.6 Workflow Summary

In summary, the full PHACTn framework proceeds through the following stages:

- **Nucleotide ancestral reconstruction (A):** Starting from a multiple sequence alignment and a phylogenetic likelihood tree, ancestral reconstruction is performed to infer nucleotide probability distributions at all internal nodes. The nucleotide ancestral states alongside a branch-optimized tree are produced.
- **Binarization and binary ancestral reconstruction (B):** The same alignment is encoded into a binary format (gap = 0, character = 1). This binary alignment is then used as input for a second two-character ancestral reconstruction. This step uses the branch-optimized tree produced in step A, with its topology held fixed, to infer indel ancestral states that capture the gap history across the phylogeny.
- **Gap-aware weighting of ancestral states (C):** The nucleotide ancestral states from step A and the indel ancestral states from step B are combined at each node by multiplying their probabilities.
- **Tree traversal and raw score accumulation (D):** Using the gap-aware ancestral states (step C) and multiple sequence alignment, the algorithm traverses the branch-optimized tree from step A, starting from the query species. At each edge, only positive probability differences are recorded and weighted inversely by phylogenetic distance. These weighted differences are then summed to generate a raw tolerance score for each nucleotide.
- **Normalization (E):** Raw tolerance scores are normalized by the total number of nodes in the tree to remove bias introduced by tree size, with an optional second normalization by the observed query nucleotide frequency to capture positional diversity.
- **Score transformation (F):** Normalized scores are mapped to a continuous [0, 1] scale via a log-based transformation, where values approaching 1 indicate likely tolerated variants and values approaching 0 indicate likely functionally disruptive variants.
- **Weighted gap frequency calculation (G):** With alignment and branch-optimized tree (step A), position-level weighted gap frequency is computed across all leaves using a distance-based weighting scheme.
- **Gap correction and final scoring (H):** The transformed PHACTn scores (step F) and the weighted gap frequency scores (step G) are integrated through a sigmoid-based correction term, producing the final PHACTn scores that incorporate both node-level and position-level gap evidence.

### 4.7 Validation Set

Disease-related variants obtained from the ClinVar [45] database in September 2025. To extract a high-confidence set of germline pathogenic single-nucleotide variants (SNVs), we only obtained variants labeled as Pathogenic/Likely Pathogenic, and we applied quality filtering based on review status: conflicting interpretations, lacking review criteria, or missing assertions were excluded. For the benign set, we processed gnomAD (v.4.1) [46] with the assumption that variants common in the population must have been selected against. Only variants with a genome allele frequency greater than 0.01 that also passed gnomAD’s internal quality controls were obtained. We merged these two-datasets to form the CG validation set, excluding any variants with conflicting significance labels. In total, 154k pathogenic and 12.8 million benign variants across the human reference genome (GRCh38/hg38) were obtained. Because of computational cost, to reduce the number of benigns, they were randomly downsampled from the same 1MB bins as the pathogenics to prevent genomic location bias. In the end, obtained benign-to-pathogenic ratio was 1:1 (~154k benign and ~154k pathogenic).

TraitGym [29] dataset obtained from the HuggingFace (songlab/TraitGym/mendelian traits). For a small number of variants, variant effect prediction could not be performed because all non-human species had gaps at the corresponding positions within the extracted window. These variants were excluded from the analysis. Since TraitGym applies a 1:9 matching scheme (one positive variant matched to nine controls), the matched counterparts of excluded variants were also removed.

### 4.8 Evaluation Metrics

The model was evaluated using five complementary metrics: area under the ROC curve (AUROC), area under the precision-recall curve (AUPR), F1 score, balanced accuracy, and Matthews correlation coefficient (MCC). Since F1 score, balanced accuracy, and MCC require a fixed classification threshold, optimal thresholds were determined by maximizing the harmonic mean of sensitivity and specificity [82]. Throughout our analysis, pathogenic variants were treated as the positive class and benign variants as the negative class.

### 4.9 Benchmark Tools

PHACTn was compared with evolutionary conservation scores (GERP [32], PhyloP [30], Phast-Cons [31]), genomic language models (DNABERT-2 [26], Nucleotide Transformer [83], Evo2 [84], AIDO.DNA [85], HyenaDNA [50]), PhyloGPN [86] and GPN-MSA [33], all of which are widely used for genome-wide variant effect prediction. Tools that use supervised machine learning for prediction were excluded from the comparison because variants present in their training sets might also appear in our benchmark dataset, leading to artificially optimistic performance estimates and preventing a fair comparison [23]. Furthermore, tools specialized for restricted genomic tasks, such as those exclusively evaluating protein-coding regions, splice sites, or specific cell-type regulatory tracks, were also omitted. Because PHACTn evaluates evolutionary constraint across the entire genome regardless of functional annotation, restricting the benchmark to genome-wide tools ensures a direct and equitable comparison. Since PHACTn is a training-free method, excluding supervised approaches and highly specialized allows for a more principled assessment of the predictive power of evolutionary signals.

**HyenaDNA [50]:** Variant effects were scored using HyenaDNA, with hyenadna-medium-160k-seqlen-hf model. For each SNV, a 131,072 bp window centered on the variant position was extracted and used to construct reference and alternative sequences. A log-likelihood ratio (LLR) score was computed between the ALT and REF alleles, averaged across both the forward and reverse complement strands to remove directional bias.

**Nucleotide Transformer v3 [83]:** Variant effect scores were calculated utilizing the NTv3 650M parameter pre-trained model (NTv3 650M pre; InstaDeep). A 131,072 bp genomic window centered on the variant position was extracted from the hg38 reference genome. The variant position was masked and the LLR score was computed as the difference in logits (log-unnormalized likelihoods) between the alternative and reference alleles at the masked site.

**AIDO.DNA [85]:** Variant pathogenicity scores were computed using the log-likelihood ratio between the reference and variant sequences at the mutation position, with a 1 kbp window centered on the variant, following the zero-shot scoring protocol described in the AIDO.DNA tutorial.

**Evo2 [84]:** Variants were scored using Evo2 (evo2 7b base), a 7-billion-parameter autoregressive genomic language model. For each SNV, an 8,192 bp window was extracted from the hg38 reference genome centered on the variant position, consistent with the model’s context length. Two allele-specific sequences were constructed per variant: a reference sequence containing the hg38 allele at the center position, and an alternate sequence with the variant allele substituted at the same position. Each sequence was independently scored using the model’s score sequences function, which computes the mean log-likelihood per nucleotide. A delta score was computed as the difference between the alternate and reference sequence log-likelihoods (ΔScore = Var Score Ref Score).

**DNABERT-2 [26]:** DNABERT2–117M was used, and for each SNV, reference and alternate sequences were constructed by extracting a 1,024 bp window from the hg38. Each sequence was tokenized using the DNABERT-2 BPE tokenizer with a maximum token limit of 512 and then passed independently through the model. We extracted CLS token and we measured variant effect as the L2 (Euclidean) distance between reference and alternate sequence embeddings.

**PhyloGPN [86]:** The prediction scores for PhyloGPN were obtained using the code available in the model’s official GitHub repository. For each single-nucleotide variant, a 481 bp sequencing window was generated from the hg38 reference genome. Sequences were processed in batches using the PhyloGPN model, and a log-likelihood ratio score was calculated for each variant as the difference between the ALT and REF nucleotide logits at the central location.

**GPN-MSA [33]:** Precomputed GPN-MSA scores were obtained from HuggingFace (https://huggingface.co/datasets/songlab/gpn-msa-hg38-scores/resolve/main/scores.tsv.bgz).

**PhyloP [30]:** Precomputed phyloP scores (100way, 470way, 30way, 447way on hg38) were obtained from the UCSC Genome Browser.

**PhastCons [31]:** Precomputed phastCons scores (100way, 470way, 30way, on hg38) were obtained from the UCSC Genome Browser.

**GERP [32]:** Precomputed GERP conservation scores were obtained from ENSEMBL (https://ftp.ensembl.org/pub/current compara/conservation scores/92 mammals. gerp conservation score).

### 4.10 Computational Cost and Runtime

All PHACTn analyses were automated using Snakemake [87] on a high-performance computing cluster, enabling reproducible and scalable execution across genomic windows. As a computational benchmark, 100 genomic windows, each 10,000 bp, were processed through the full pipeline with up to 50 parallel jobs. Overall consumption was approximately 27 CPU-hours across 800 jobs with a peak memory of 1.1 GB per job, demonstrating that PHACTn operates efficiently on standard CPU nodes.

## 5 Declarations

### 5.1 Data and resource availability

The source code for PHACTn, including installation guidelines, execution instructions, and the core scoring algorithms, will be publicly available on GitHub at https://github.com/ CompGenomeLab/PHACTn upon acceptance. The precomputed whole-genome scores generated in this study will also be made publicly available upon manuscript acceptance at Zenodo.

All other analyzed datasets are from public sources: ClinVar, gnomAD v4.1, the UCSC 470-way mammalian alignment and GRCh38 reference (UCSC Genome Browser), TraitGym Mendelian traits (HuggingFace, songlab/TraitGym), and precomputed GPN-MSA, phyloP, phastCons, and GERP scores as described in Methods.

### 5.2 Competing interests

The authors declare that they have no competing interests.

## 5.3 Acknowledgements

This work is part of the PHACT-miRBind project, supported by Xjenza Malta and the Scientific and Technological Research Council of Türkiye (TÜBİTAK, Grant No. 124N485, awarded to O.A.) through the MCST/TÜBİTAK Joint Call for Research and Innovation in Artificial Intelligence in Health.

The numerical calculations reported in this paper were partially performed at TÜBİTAK ULAKBIM, High Performance and Grid Computing Center (TRUBA resources).

---

**Fig. S1:**
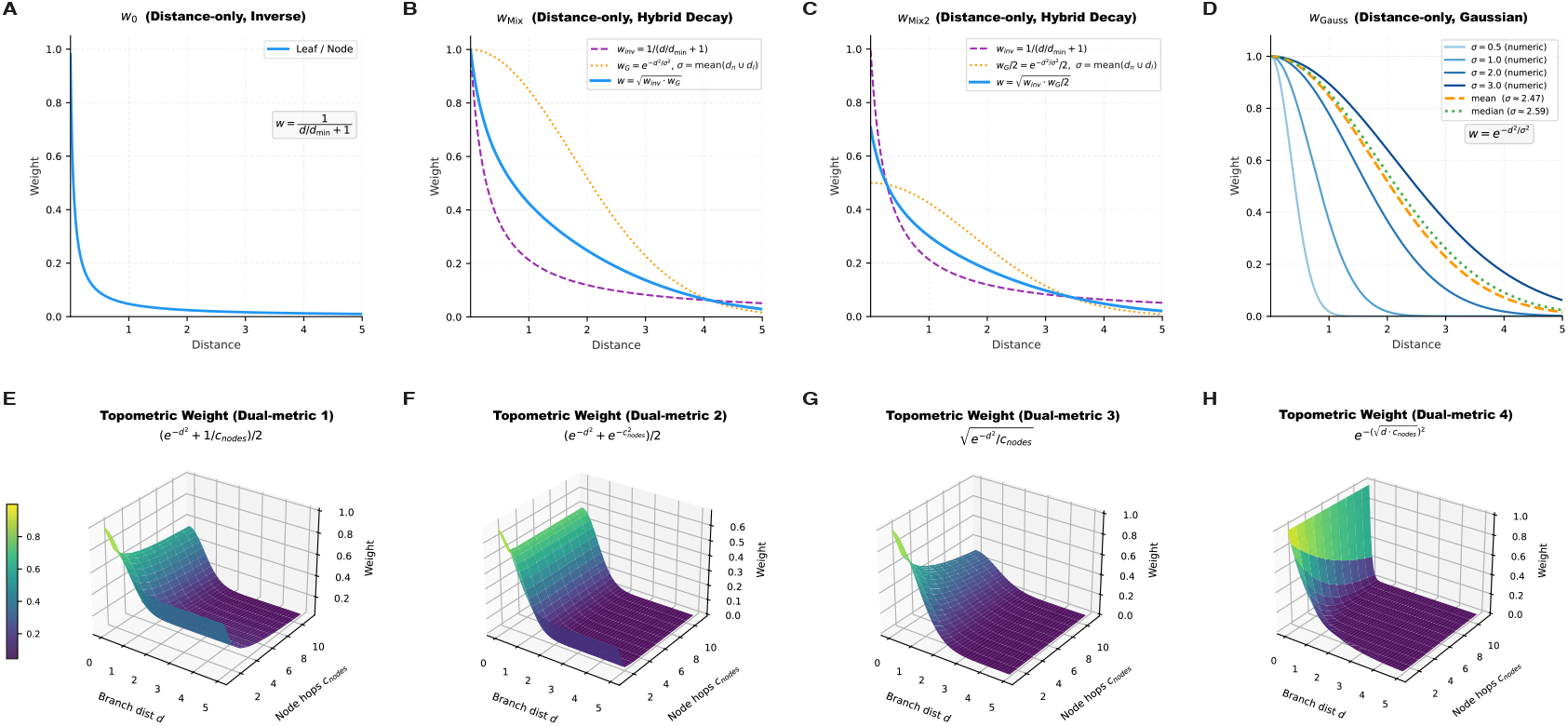
Weight functions used in PHACTn score accumulation, grouped by the information they incorporate. **(A–D)** Distance-only schemes, in which weight depends solely on branch length *d* to the human leaf. **(A)** *w*_0_: inverse decay normalised by the minimum node distance, *w* = 1*/*(*d/d*_min_ + 1). **(B)** *w*_Mix_: geometric mean of the inverse and a Gaussian component, 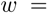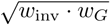, where 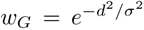 with *σ* = mean(*d_n_ ∪ d_l_*). **(C)** *w*_Mix2_: same geometric mean construction but with the Gaussian amplitude halved, 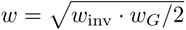, yielding a moderately faster decay than w_Mix_. **(D)** *w*_Gauss_: pure Gaussian decay 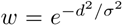 shown for six σ choices: fixed values σ ∈ {0.5, 1.0, 2.0, 3.0}, the tree-adaptive mean (*σ ≈* 2.47 in this example) and median (*σ ≈* 2.59 in this example) of all branch lengths. **(E–H)** Topometric (dual-metric) schemes, in which weight depends jointly on branch length *d* and the number of ancestral nodes *c*_nodes_ on the path between the current node and the human leaf; surfaces are shown over the domain *d ∈* [0, 5], *c*_nodes_ *∈* [1, 11]. **(E)** Dual-metric 1: arithmetic mean of a unit Gaussian and an inverse hop term, 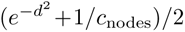. **(F)** Dual-metric 2: arithmetic mean of two Gaussian terms, 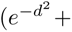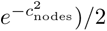. **(G)** Dual-metric 3: geometric mean of the unit Gaussian and the inverse hop term, 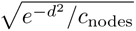. **(H)** Dual-metric 4: coupled exponential penalising jointly large distance and hop count, 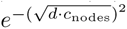 . Colour encodes weight magnitude in panels E–H (yellow: high, purple: low).

**Fig. S2:**
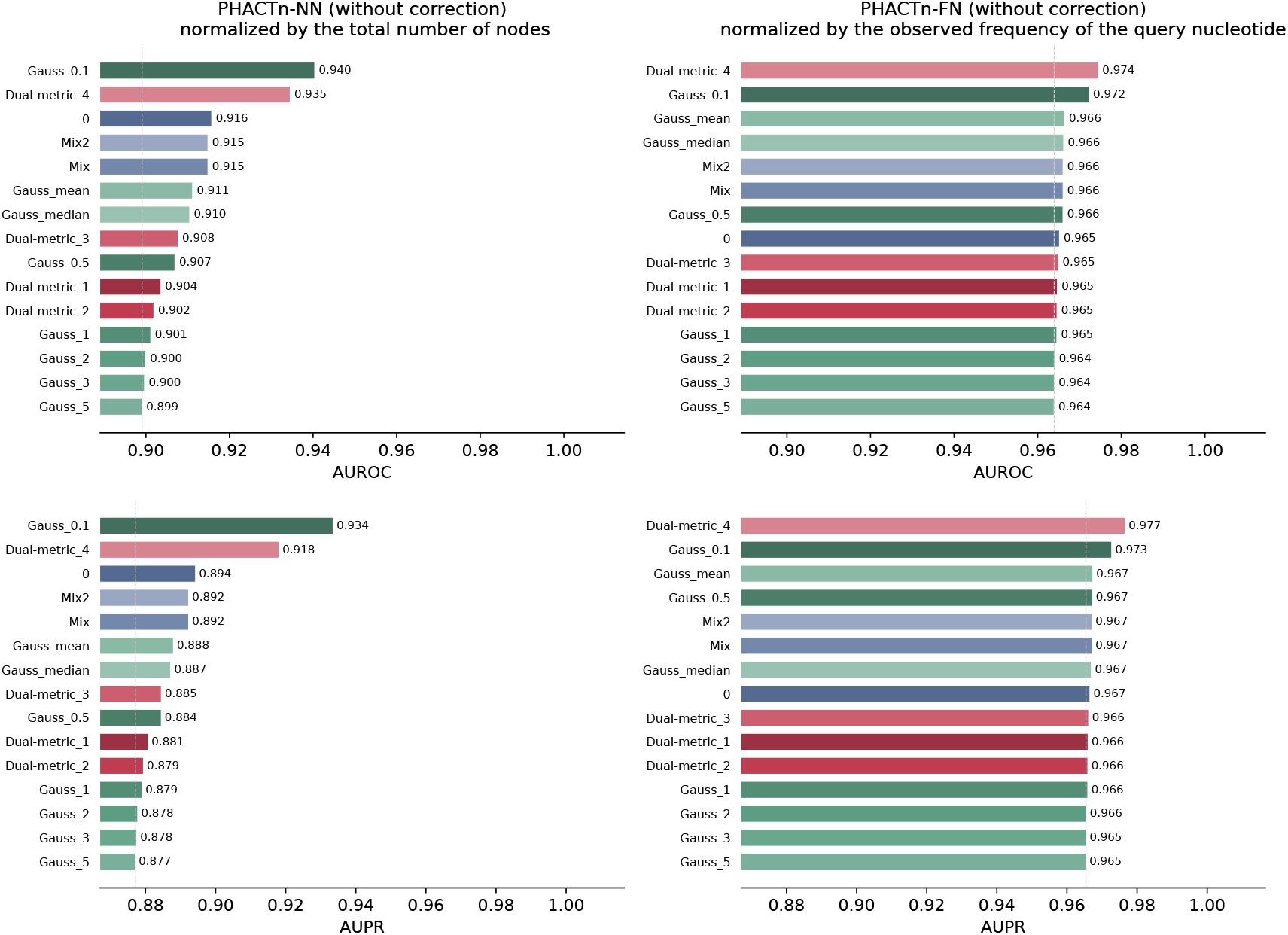
Performance comparison of weighting functions using base PHACTn (without gap correction) on the ClinVar-gnomAD variant set. AUROC (top) and AUPR (bottom) are shown for all evaluated weighting functions under two normalization strategies: normalization by the total number of nodes (PHACTn-NN, left) and normalization by the observed frequency of the query nucleotide (PHACTn-FN, right).

**Fig. S3:**
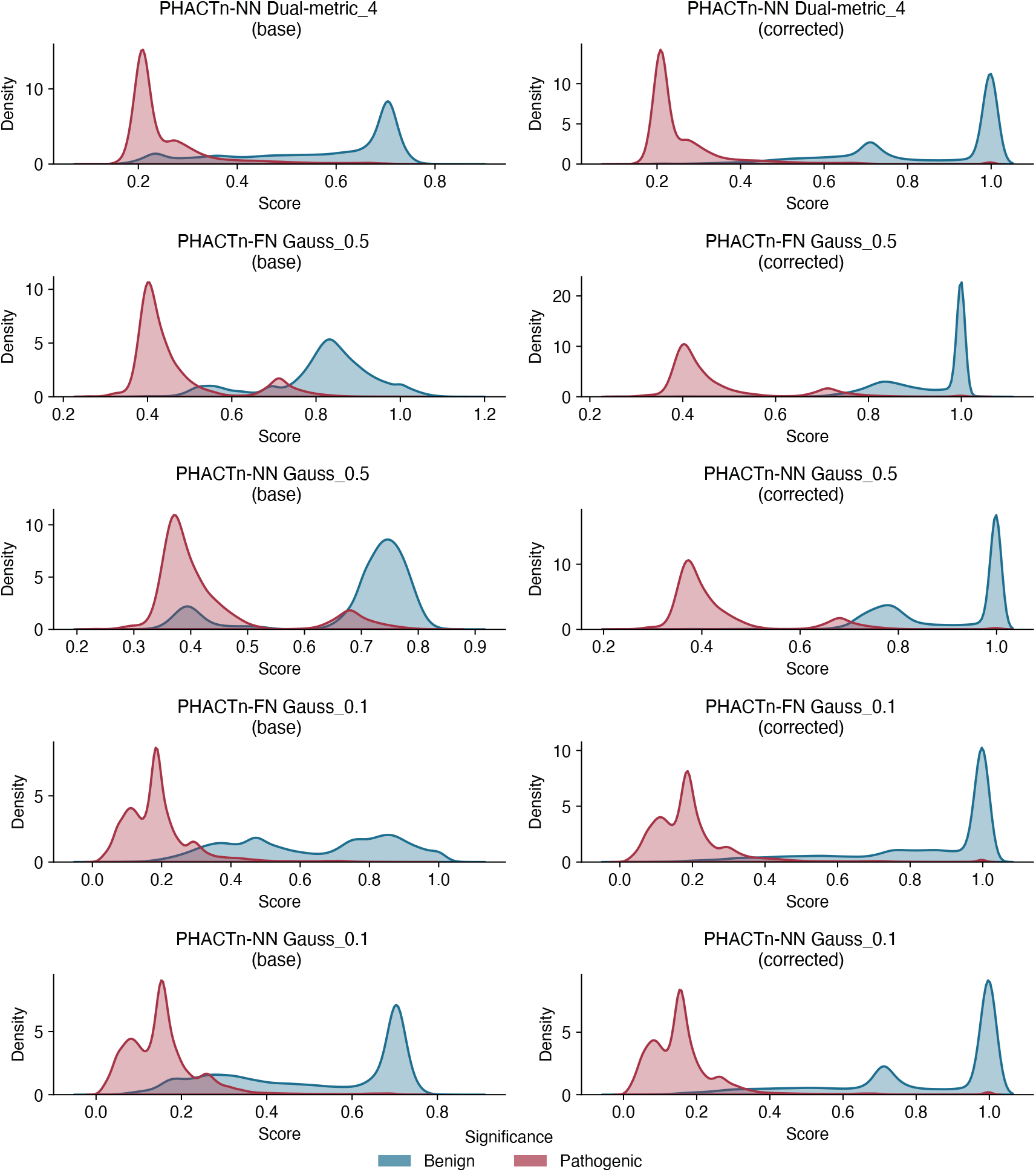
Score distributions for benign (blue) and pathogenic (pink) variants before and after gap correction, across three PHACTn model variants. Correction consistently shifts benign scores toward 1.0, sharpening separation between the two classes.

**Fig. S4:**
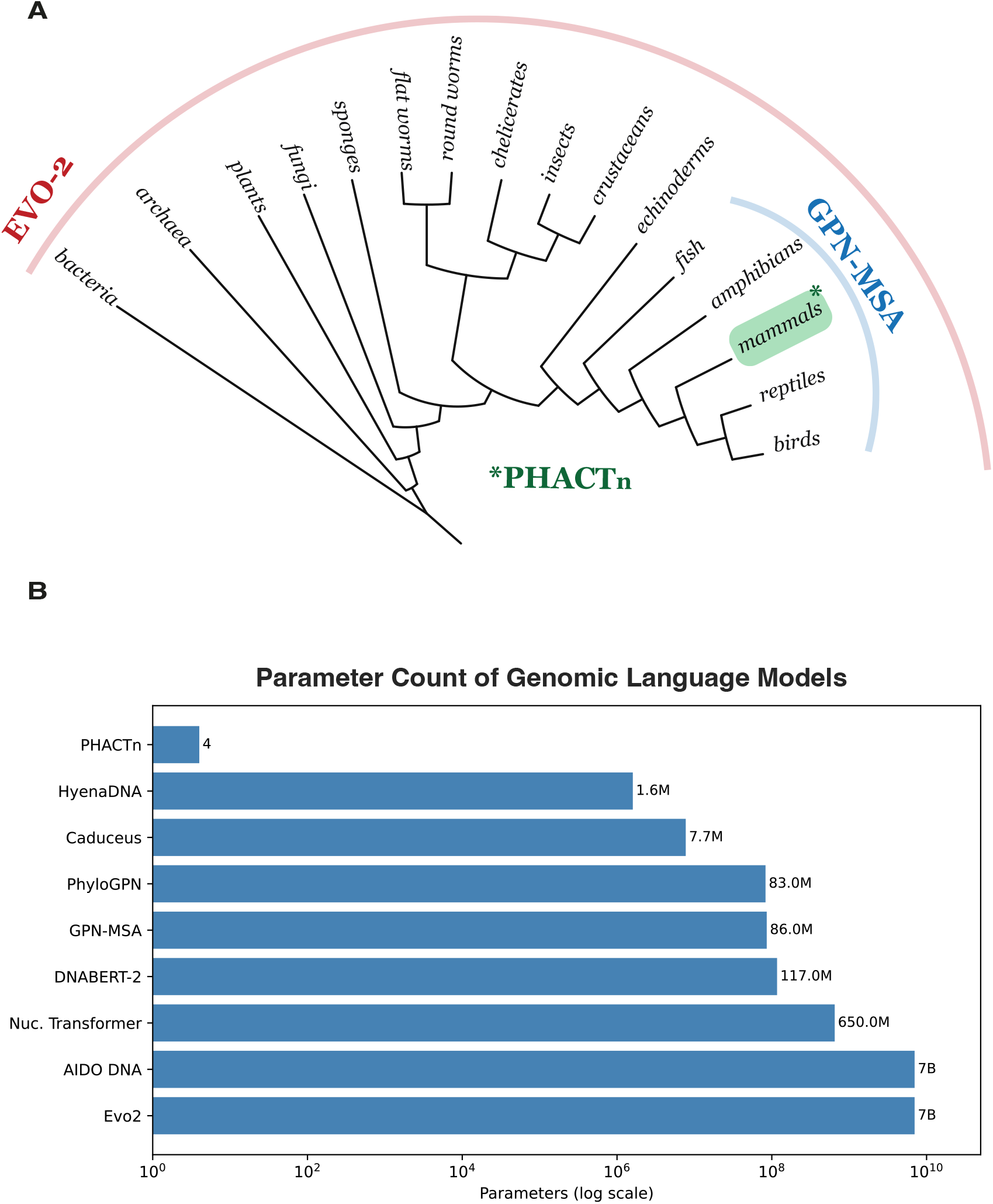
(a) Phylogenetic scope of PHACTn, GPN-MSA, and Evo-2. PHACTn (green) operates exclusively on the 470-way mammalian alignment; GPN-MSA (blue) spans the full vertebrate clade including a 100-vertebrate whole-genome alignment; Evo-2 (red) was pre-trained on sequences from all domains of life. (b) Parameter counts of PHACTn and DNA foundation models on a log scale. PHACTn uses 4 interpretable parameters, spanning more than nine orders of magnitude fewer parameters than the largest models.

**Fig. S5:**
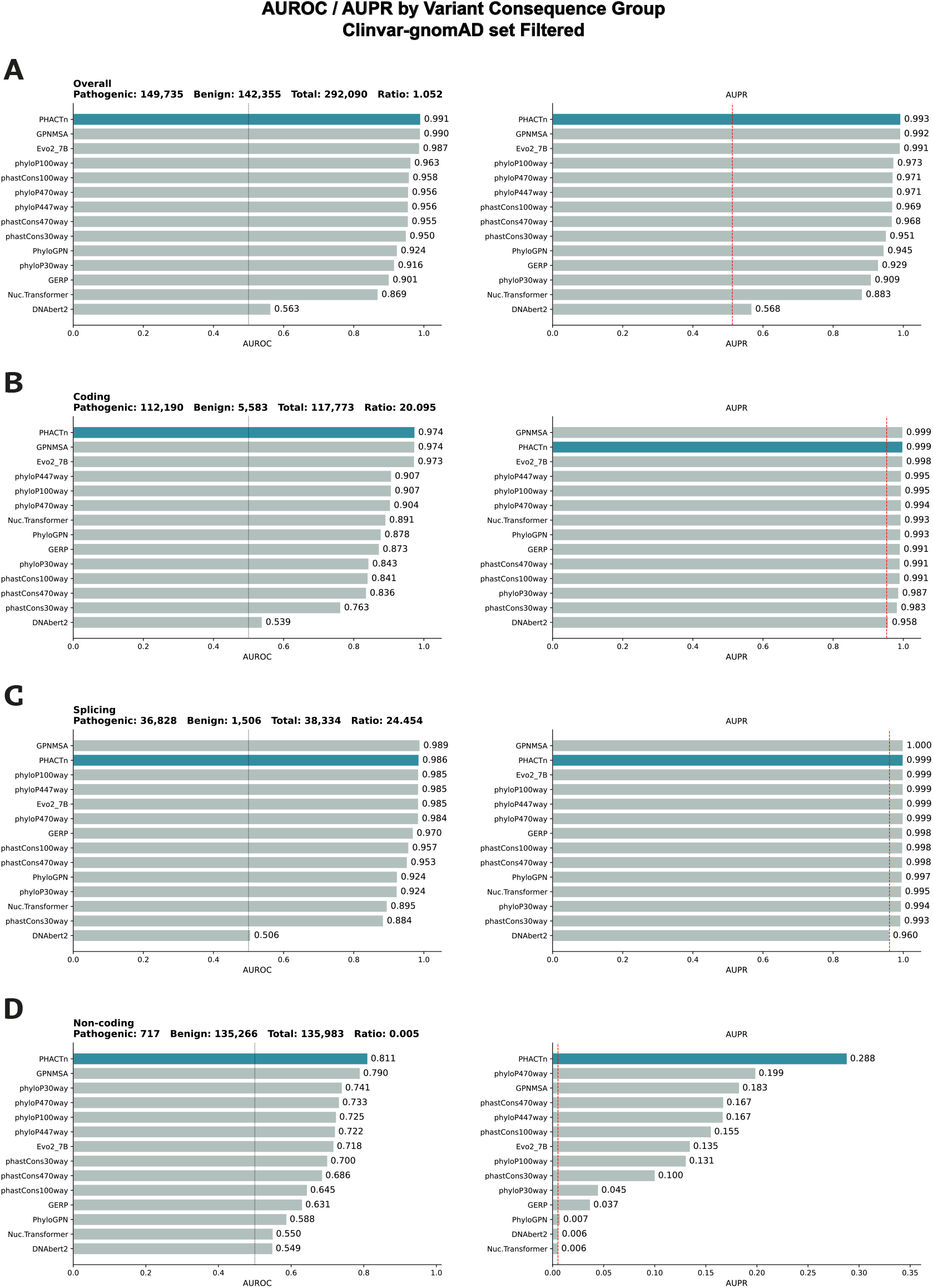
Performances on the ClinVar–gnomAD variant filtered set on problematic cases (alternative frequency = 0, gap frequency ≤ 0.3, and benign) AUROC and AUPR are shown for **(a)** overall, **(b)** coding, **(c)** splicing, **(d)** non-coding.

## Notes

### Competing Interest Statement

The authors have declared no competing interest.

### Summary of Updates

The Acknowledgments section has been updated to include details on funding and grant support for the project.

